# Genomic Evidence of Fisheries Induced Evolution in Eastern Baltic cod

**DOI:** 10.1101/2024.06.27.601002

**Authors:** Kwi Young Han, Reid S. Brennan, Christopher T. Monk, Sissel Jentoft, Cecilia Helmerson, Jan Dierking, Karin Hüssy, Érika Endo Kokubun, Janina Fuss, Ben Krause- Kyora, Tonny B. Thomsen, Benjamin D. Heredia, Thorsten B.H. Reusch

## Abstract

Humans have become one of the greatest evolutionary forces, and their perturbations are expected to elicit strong evolutionary responses. Accordingly, during (size) selective overharvesting of wild populations, marked phenotypic changes have been documented, while the evolutionary basis is often unresolved. Time-series collections combined with genomic tools present unique opportunities to study how evolutionary changes are manifested at the genome-wide level. Here, we take advantage of a unique temporal dataset from the overexploited Eastern Baltic cod (*Gadus morhua*) population that exhibited a 48% decrease in asymptotic body length over the last 25 years. A genome-wide association study revealed pronounced peaks of outliers linked to growth performance. The contributing loci showed signals of directional selection with significantly high autocovariance in the allele frequency and excessive intersections with regions of high F_st_ as well as genes relevant to growth and reproduction. Moreover, pattern of directional selection for ancestral haplotype of the well-known chromosomal inversions in Atlantic cod (on linkage group 12) was observed, while the double crossover (∼1Mb) enharbouring the vitellogenin genes within this region showed signs of drift or balancing selection. Our results demonstrate evident response of the genome over a relatively short time frame and further underscore implications for fisheries management and conservation policy regarding the adaptive potential of marine populations.

## INTRODUCTION

Human beings play a significant ecological and evolutionary role as they manipulate and disrupt environments and organisms by habitat alteration, pollution, climate change and harvesting (Palumbi 2001). This impact extends beyond a population’s distribution and its relevant ecological landscape of one time point and influences future generations by exerting strong selective pressures (Vitousek et al. 1997; Palumbi 2001; Hendry, Gotanda, and Svensson 2017). Rapid evolutionary changes caused by anthropogenic pressures, e.g., overfishing, pose special challenges in detecting induced selection processes, as the changes usually span a relatively short time frame insufficient for a conventional sweep-like pattern causing the complete fixation of focal alleles. Here, historical time-series samples provide a special lens to the past in detecting evolution in action by enabling direct access to allele frequency changes in genomic data (Franssen, Kofler, and Schlötterer 2017). In the context of fisheries induced evolution, one of the strongest human perturbations caused by size selectivity or added mortality onto a fish population, so far, the only compelling evidence for genome-level responses to overfishing comes from a 40 years of annual time series data of Atlantic salmon. A clear decrease in age at maturity in Atlantic salmon was accompanied by directional change in the allele frequency of *vgll*3 gene (Erkinaro et al. 2019; Czorlich et al. 2018), a large effect locus explaining 39% of the phenotypic variation (Barson et al. 2015; Ayllon et al. 2015), which was significantly correlated with fishing pressure for the target species as well as a food species in salmon aquacultures (Czorlich et al. 2022). However, as most traits under fishing induced selection, including life history traits like growth rate, have a polygenic basis with a large number of small effect loci, challenges remain in both the identification of the contributing loci and the detection of subtle changes in frequency of the loci (see (Reid, Star, and Pinsky 2023; Pinsky et al. 2021)).

Eastern Baltic cod (EBC) is an Atlantic cod (*Gadus morhua*) population residing in the central Baltic Sea, with the last remaining spawning ground being the Bornholm Basin (ICES 2022). The population diverged from other Atlantic cod populations 7-8 thousand years ago when the Baltic Sea with its current salinity regime emerged after a series of postglacial tectonic shifts in combination with sea-level changes (Matschiner et al. 2022; Schmölcke et al. 2006; Martínez-García et al. 2021). Currently, it is biologically and genetically differentiated from all other ecotypes, e.g. western Baltic (WBC) and North Sea cod and adapted to the peculiar Baltic environment and experiences low salinities, high pCO2, prevalent hypoxia, and inconsistent and highly variable seasonal patterns of temperature, salinity and oxygen contents (Reusch et al. 2018; Zillén et al. 2008; Stockmayer and Lehmann 2023). These fluctuating environmental conditions have contributed to the indistinguishable pattern in the otolith rings for age readings, compromising the age-related data for stock assessments of EBC (Heimbrand et al. 2020). At present, EBC is isolated from neighbouring WBC in the absence of genetic inflow (Paul R. Berg et al. 2015; Hemmer-Hansen et al. 2019), even though some limited hybridization occurred historically, at high population abundance (Helmerson et al. 2023).

EBC plays a major role not only ecologically as a key predator species in the food web, particularly notable in the Baltic’s uniquely low biodiversity (Ojaveer et al. 2010), but also economically as it has been fished recreationally and was the largest target species for commercial fisheries with an annual catch of up to 400,000 tons in the mid-1980s (ICES 2022). However, overfishing and size-selective fishing continued with excessively high total allowable catch resulting in fishing mortality typically 2-3 times higher than the maximum sustainable yield (MSY) (Birgersson 2022; ICES 2019; Eero et al. 2011; Zeller et al. 2011). Since the mid-1990s, multiple aspects of the EBC population have been deteriorating and have recently reached the unprecedented lowest point in their state since the 1950s (Birgersson 2022; Eero et al. 2023). The spawning stock biomass (fish sized over 35cm) has declined sharply in recent years, together with recruitment and loss of two major spawning grounds (Cardinale and Svedäng 2011; Köster et al. 2017). Higher mortality on older individuals can lead to size truncation, growth retardation, and worsened condition (weight-at-length) (Eero et al. 2023; Möllmann et al. 2009; Svedäng and Hornborg 2014; 2017). The size at first maturity and condition of the fish marked the lowest value of L50 (length at 50% of population reaches maturity) under 20 cm in recent years (Eero et al. 2015; ICES 2021; Mion et al. 2021; Svedäng and Hornborg 2017). A complete collapse of the stock has resulted in a ban on targeted fishing on EBC since 2019 (ban renewed for 2024) but the condition of the population has not been able to recover to a healthy status so far.

Despite the prominent changes in body length in the EBC, the genetic basis of the change, thus the evolutionary consequences of overfishing, has not been investigated until now. Here, we investigated whether or not changes in a heritable trait under selection caused by size-selective trawling translates to a detectable response of the genomes over time. To this end, we modelled individual growth using archived otoliths and sequenced whole-genomes of the population over multiple time points in the period of 1996-2019 (referred as “temporal population” hereafter). Individuals caught in Bornholm Basin (Figure 1A) were selected to cover the full breadth of time and phenotype spectrum, by random sampling along the length distribution for each time point (a sample set called “random” hereafter), then including individuals at the both tails of the distribution (called “phenotype”). A genotype-phenotype association study (GWAS) identified pronounced peaks of outlier loci near genes linked to growth and maturity, which in turn showed signals of selection. In parallel, we found a heterogeneous pattern of selection in a large inverted region in linkage group (LG) 12. This study is, to the best of our knowledge, the first in a fully marine species to provide leads that suggest genomic changes to underlie phenotypic evolution of a polygenic trait in response to overfishing in the field. It showcases the strength of combining temporal genomics of wild population with its phenotype data and eventually guides us through connecting dots of fisheries induced evolution.

**Figure 1.**
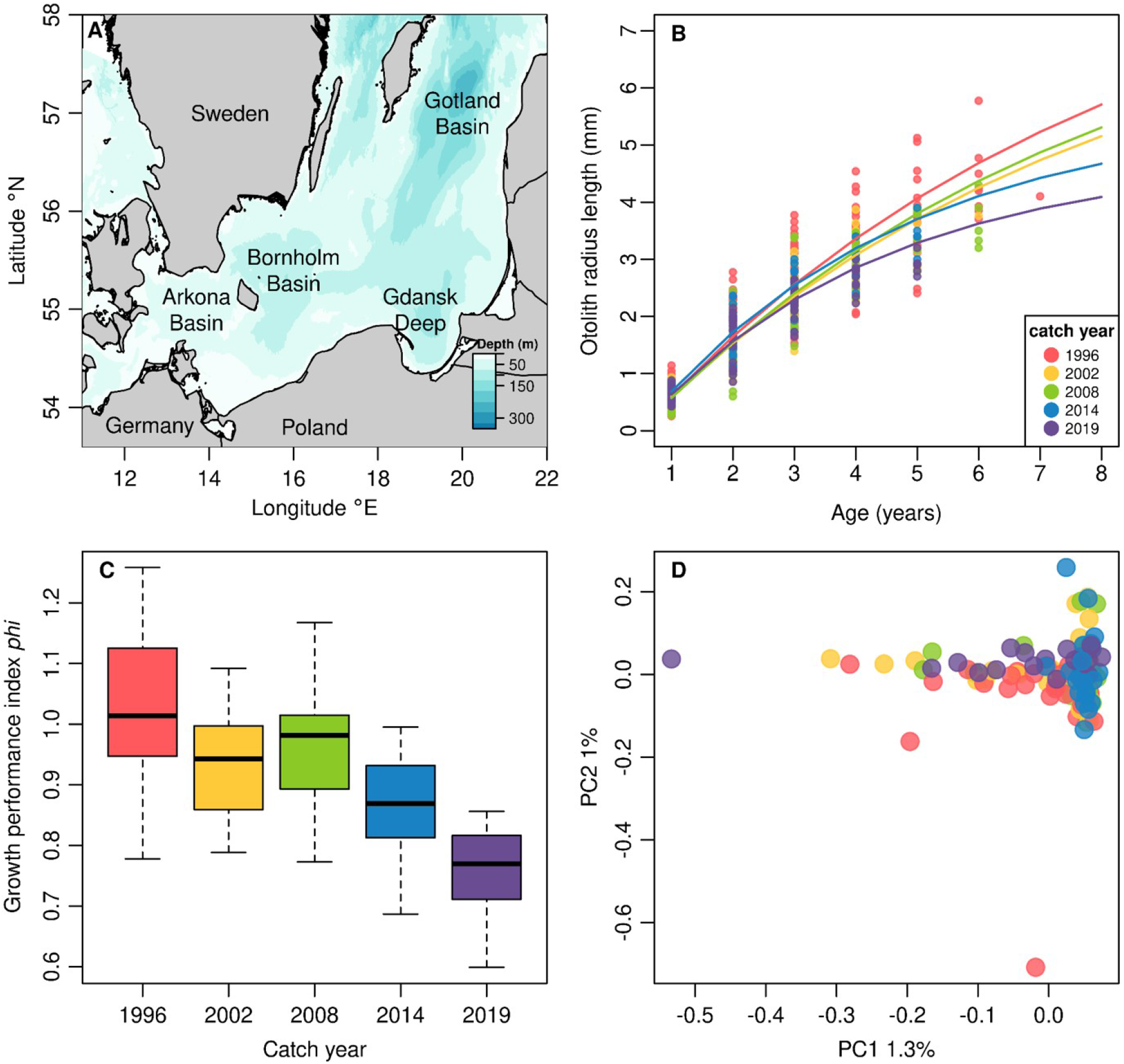
Sampling location in the Baltic Sea and population response over time. **A.** Map of the Baltic Sea showing the sampling sites in Bornholm Basin, the major spawning ground for EBC. Their spawning grounds in Gotland Basin and Gdansk Deep are not recognized as viable anymore. **B.** Estimated von Bertalanffy growth curves for each catch year. The von Bertalanffy growth curves are based on otolith readings and were plotted using estimated sets of parameters for each temporal population of “random” and “phenotype” samples. The temporal group 1996 in this figure also includes “phenotype” samples (catch year from 1996-1998) as they are treated as one temporal population in the model. Each point depicts observed otolith radius to chemical annuli at age coloured based on the individuals’ catch year. **C.** Boxplots of individual growth performance, Φ, calculated using estimated individual von Bertalanffy growth parameters (*L∞* and *k)* over time. Colour codes are based on individuals’ catch years as in the legend in panel B. **D**. Principal component analysis of 115 “random” samples. A set of SNPs were pruned based on linkage disequilibrium and removed of sites within the inversions in LG2, 7, and 12. PC1 explains 1.3%, and 1% for PC2, of all variations in the genotypes. Each individual is coded in colour according to the catch years as in legend in panel B.

## RESULTS

### Temporal Changes in Growth Rates

To demonstrate a phenotypic change under size-selective fishing pressure over the last 25 years (1996-2019), we focused on individual growth rates as the key heritable trait. We first aged archived otolith samples of 152 EBC individuals from Bornholm Basin using a novel method of biochemical reading as age information of EBC recorded through a conventional method has been unreliable (Hüssy et al. 2021). The oldest fish was a 7-year-old caught in 1996 while individuals as old as 5 years old could be sampled in more recent years (2014 and 2019) (Table S1). Using the distance from the core to each chemical annulus, the estimated otolith radii, von Bertalanffy growth parameters were estimated for each fish individual and each temporal population (von Bertalanffy 1957); Table S2). Fish in 1996 grew to reach a larger terminal maximum size, and had a smaller Brody growth coefficient k, meaning they took longer to approach their terminal length than fish from recent years. The median of estimated individual length at infinity, *L∞*, decreased by 48% from 1996 to 2019, with a small inconsistency in 2008 (Figure 1B and S1A). Remarkably, this translates to a maximal fish length (*L∞*) decrease from 1150 mm in 1996 to 539 mm in 2019 when back-calculating fish body length from otolith radii. Accordingly, growth coefficient *k* increases over the period with the same trend in 2008 in both group parameters and individual parameters (Figure S1B). A growth performance index (Φ) for each fish of different years, which summarises the growth (Moreau, Bambino, and Pauly 1986), showed a consistent decrease in time (Figure 1C). Additionally, the otolith radii at age 1 for all fish were back-calculated to body length and compared to examine any deviation in the juvenile growth of EBC in temporal trend (Figure S1C). Although mean distances to the first-year radii do not differ, the variance of the radii significantly reduced over time (Bartlett’s test for variance, p-value = 0.02868), indicating truncated phenotypic diversity in juvenile growth. Here, the condition of individual fish at catch (relative condition factor (Le Cren 1951)) showed statistically different population mean only for 2002 (Figure S2A). When tested for correlation, individuals’ condition did not predict either the growth parameters, *L∞* and *k*, or Φ (r = −0.03 (p > 0.05), r = 0.09 (p > 0.05), and r = 0.09 (p > 0.05) respectively). (Figure S2B- D). Overall, this supports that the population has shifted to grow slower and reach smaller size when older during the study period of heavy fishing pressure.

### Genome-wide Temporal differentiation

In order to investigate any temporal differentiation of EBC which might potentially correspond to the phenotypic change, we subjected a set of 5,847,389 SNPs (MAF > 0.005) identified for 115 “random” samples to population summary statistics. First, a principal component analysis (PCA) using SNPs outside previously reported large chromosomal inversions revealed a panmictic population structure among time points (Figure 1D). The variances explained by PC1 and PC2 were relatively small (1.26 and 1.03%) while the loadings for each PC were well distributed along the whole genome. Second, we applied a temporal covariance analysis developed by Buffalo and Coop (Buffalo and Coop 2019; 2020) to test genome-wide pattern of selection signature. The pairwise autocovariance of allele frequency changes in all time windows showed a pattern that resembled that of a simulated neutral scenario (Figure S3 and S4). The observed temporal autocovariance values from the samples were within the distribution of the expected under drift (p > 0.05 for all paired autocovariances), which shows a lack of genome-wide selection signal (Figure S5). Lastly, genome-wide nucleotide diversity (**π**) and absolute divergence between populations (dxy), calculated for 50 kb - windows varied only little among years (Figure S6). As expected, windowed **π** and dxy varied along linkage groups depending on differences in recombination rate along the chromosome, e.g. centromere regions featuring less recombination (Sardell and Kirkpatrick 2020; Tigano et al. 2021). Some divergence was observed at the beginning of LG2 and in the central section of LG7, which were most likely caused by the varying frequency of the inverted regions. Overall pattern shows comparable genome-wide **π** (ranging from lowest value of 0.0071 for 1996 to highest value of 0.0077 for 2008) and consistent slight increase in dxy values as the sampling points are more distant, which indicates drift over time.

### Genotype-Phenotype Association and Selection of SNPs linked to growth

In the lack of genome-wide signal of selection, we sought to identify loci under directional selection by a genome-wide association (GWA) analysis using individual growth performance index (Φ) as a phenotype and 679,584 biallelic SNPs (MAF > 0.05). Three regions of the genome were clear outlier peaks with -log10P values around 6 and most likely to be associated with growth performance (Figure 2). The distribution of moderate p-values (3 < -log10P <5) across the genome in itself shows the polygenic nature of growth as well as the methodological limitations of GWA analysis given our relatively low sample size of 152. Under a formal correction for multiple testing only a few regions remained at p = 0.05. As this study was designed as an explorative approach, an outlier status was assigned to 338 SNP loci that lie in the lowest 0.05% of the distribution of p-values.

**Figure 2.**
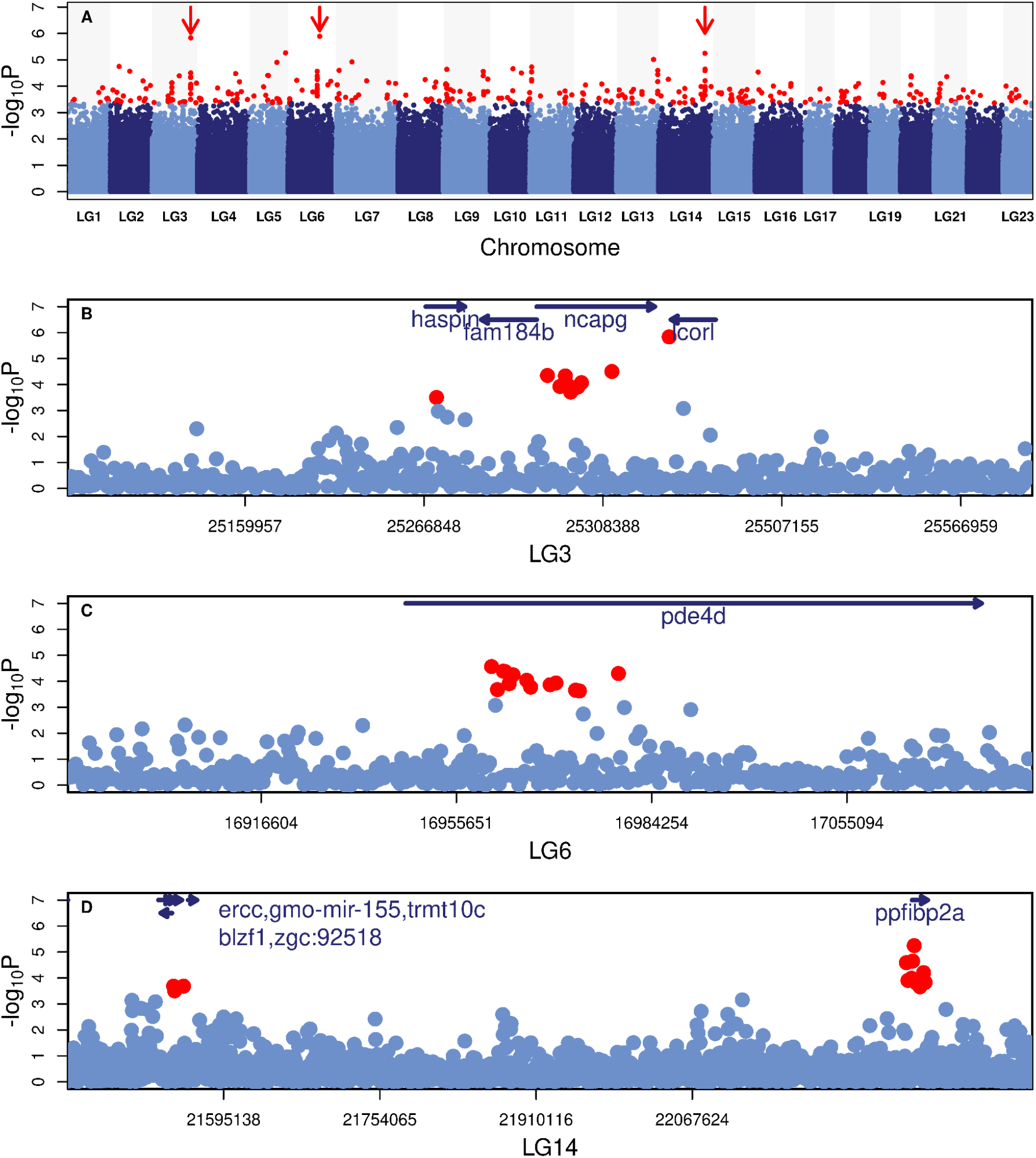
Manhattan plot of -logP values in genome-wide association (GWA) analysis. **A.** Manhattan plot of -logP values in genome-wide association (GWA) analysis. A total of 152 samples were subjected to GWA using the sequenced genotypes, 679,584 SNPs (>0.05 MAF), and estimated growth performance index Φ as phenotype. Negative log transformed Wald test p-values for each SNP were plotted along the genome. Outlier status was assigned for 338 SNPs with lowest 0.05% p-values (in red circles). The cutoff for outliers were selected based on the visual examination of this Manhattan plot, so as to include distinctive peaks with clustering outliers (marked with red arrows) and at the same time exclude spurious outliers consisting of single SNPs only. Regions marked with red arrows were zoomed in, **B.** in LG3 **C.** LG6, and **D.** LG14 and genes residing at or near (5 Kb up- and downstream) the outliers are annotated (Table 1).

**Table 1.**
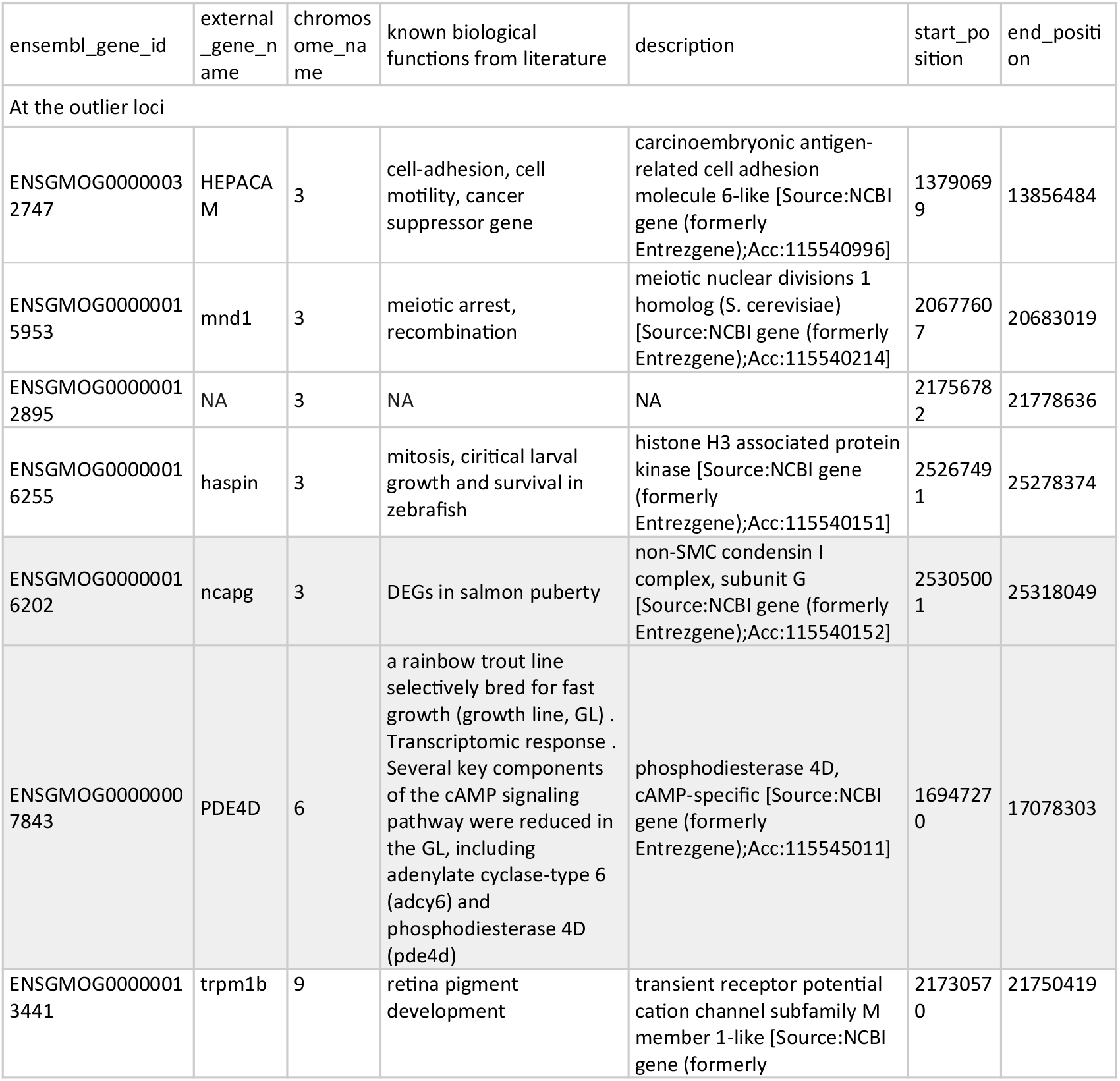
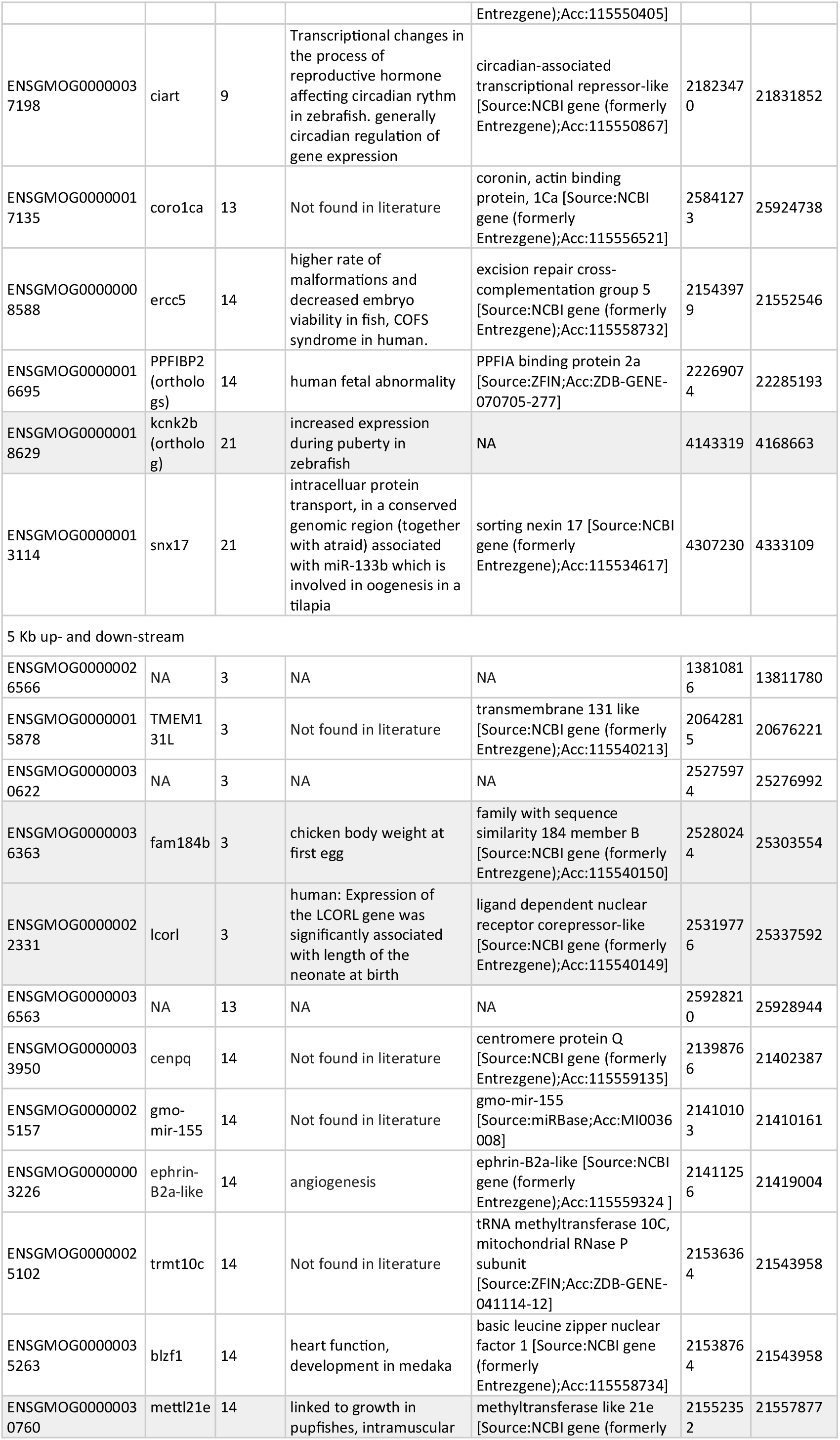
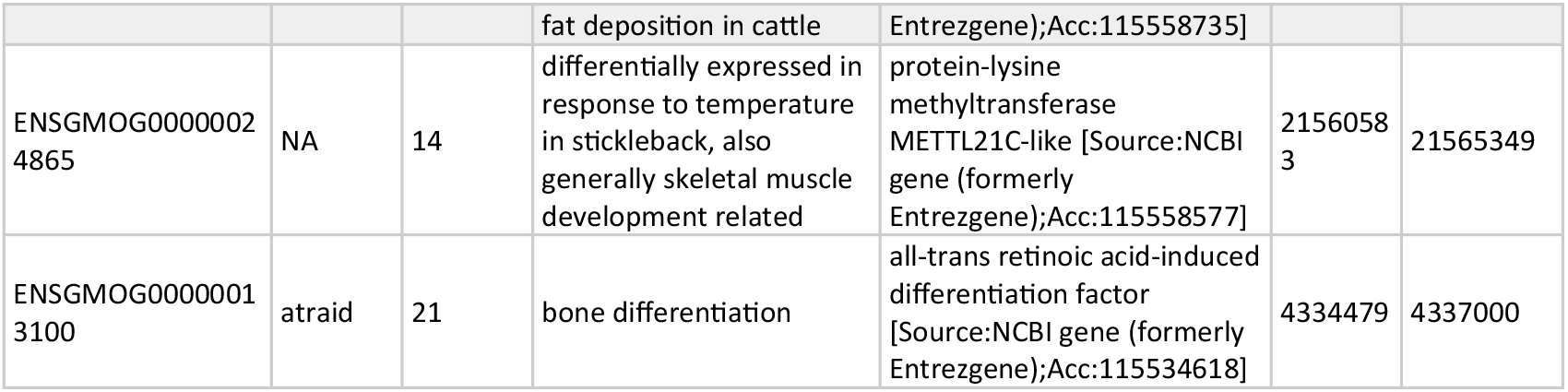
A list of genes intersecting or neighbouring with GWA outlier SNPs with its position and description. A list of genes unique in the “ensembl_gene_id” located at or 10 Kb surrounding regions of GWA outlier SNPs was subjected to a search for their functional annotations in the literature. Columns, ensembl_gene_id, chromosome_name, external_gene_name, description, start_position, and end_position, are annotations in the gadMor3.0 reference genome extracted from Ensembl database. When the external_gene_name were not provided by the database, search in the NCBI gene database or orthologs’ name were filled in (Genes, NCBI). Functional annotations of the genes, in the most relevant context to this study, “known biological functions from literature", were listed by searching the gene names with or without keywords (e.g., fish, growth, weight, maturity, and reproduction) in the literature search. When there were no search results which showed direct or indirect biological relevance in the targeted context, they were marked as “Not found in literature". Some genes were catalogued with only weak matches to orthologs in other species in the database, thus marked as “NA (Not applicable)". Rows containing genes which are most relevant to this study are highlighted in grey.

Regions with a peak of clustered outliers with flanking SNPs with low p-values were examined in depth to seek biological relevance of the SNP sites. Genes which span over 5 Kb up- and downstream of the outliers were listed as candidate genes linked to growth variations. Amongst these candidate genes, the three most evident peaks of outliers in LG3, LG6 and LG14 contained genes which were most relevant to growth or maturity from functional annotation and previous research (Figure 2B-D, Table 1): LG3 contains *ncapg*, which is differentially expressed in puberty in salmon (Crespo et al. 2019) and *fam184b*, which is associated with body weight at first egg in chicken (Fan et al. 2017). Linkage group 6 included *pde4d* gene which showed response in the transcriptome of fast growth line in a rainbow trout (Cleveland, Gao, and Leeds 2020). Finally, in linkage group 14 *mettl21e* which was linked to growth in pupfishes and intramuscular fat deposition in cattle (Fonseca et al. 2020; Patton et al. 2022).

In order to understand if the genomic regions explaining phenotypic variation were under selection through time, we calculated covariance values for the GWA outliers to observe directional change in their allele frequency. Specifically, lag-2 (i.e. cov(Δ1996-2008, Δ2002-2014) and cov(Δ2002-2014, Δ2008-2019)) and lag-3 (i.e. cov(Δ1996-2014, Δ2002-2019)) autocovariance (as illustrated in inlets of Figure S7) were calculated. Temporal covariances of allele frequency changes of 338 outlier SNPs exhibited remarkably high values of 0.00154 and 0.00187 for lag-2 and 0.00537 for lag-3 (Figure S7). Based on 1000 random permutations of covariance values of 338 SNPs sites, the observed covariances of GWA outliers markedly exceed the ranges of null-distributions (p < 0.001). This result strongly supports that the GWA outliers, highly correlated to the growth performance collectively, experienced selection and responded accordingly with a directional frequency change over time.

### Integration of the selection scan and GWAS

As a complementary approach to detect directional selection of loci linked to growth, we combined the GWA results with a selection test. An F_st_ scan on 20 Kb sliding windows across the genome was conducted comparing the temporal population of 1996 and 2019. Despite the lack of genome-wide signal of selection among temporal populations, we were able to identify regions of higher differentiation (Figure 3). While the genome-wide F_st_ value was xxx, a low value as expected for a single spatial population, some regions showed higher F_st_ values up to 0.1. When outlier windows of 5% highest p-values were assigned to intersect with GWAS outliers, 33 windows overlapped. To test the statistical significance of this overlap, a null distribution was produced with a randomization test which the observed values can be compared to. Based on 5000 random permutations, wherein 338 SNPs were randomly chosen to overlap with the outlier windows, the observed number exceeded the upper tail of the expected distribution (Figure S8). This signifies that loci associated with growth performance are predicted to reside in the regions of highest F_st_ between 1996 and 2019. Those two lines of evidence, the positive temporal covariance values of GWA outliers and their significant overlap with high F_st_ windows, strongly indicate the impact of directional selection on the genetic factors under growth variations in EBC.

**Figure 3.**
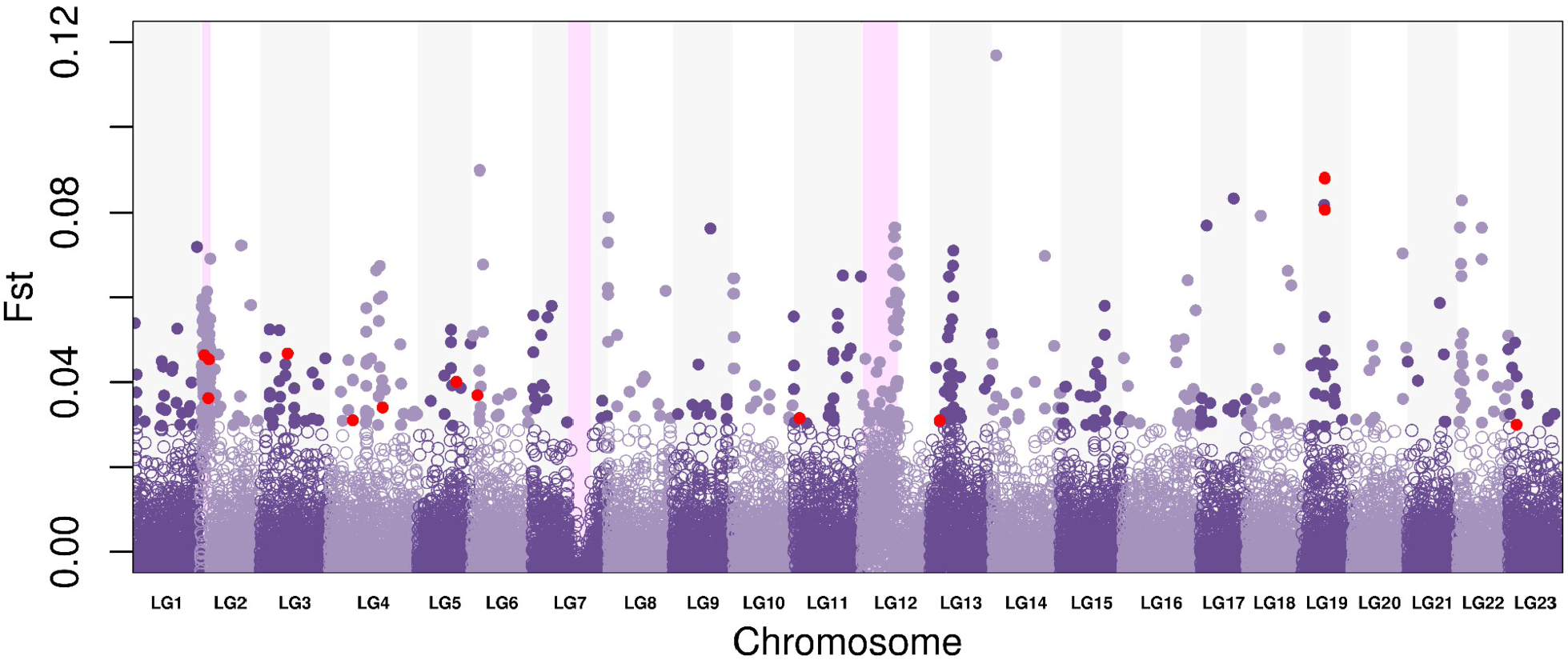
F_st_ values in 20kb windows along the genome. Pairwise F_st_ values of 1996 and 2019 in 20 Kb non-overlapping windows were calculated along the whole genome. Filled purple points indicate the highest top 5% of genome-wide F_st_ outlier windows. Regions with exceptionally high and low F_st_ values show the inverted regions in cod genome in LG2, LG7, and LG12 and were marked with pink shades. Among outlier windows, windows overlapping with GWA outliers were marked as red.

The biological significance of these overlapping regions was further explored through a gene ontology (GO) term enrichment test on overlapping F_st_ windows (Table 2). Multiple pathways involved in ultradian rhythm, water homeostasis, and protein metabolism, and meiotic cell cycle were enriched. Ultradian rhythm is important in diverse functions including growth, reproduction, and metabolism in fish (Cowan, Azpeleta, and López-Olmeda 2017; Frøland Steindal and Whitmore 2019; Sánchez-Vázquez et al. 2019; Zhdanova and Reebs 2006). Diverse metabolic processes involving amino acids were also significantly enriched, which is critical for fish growth rates (Finn and Fyhn 2010; Pelletier et al. 1994). Interestingly, folic acid deficiency in diet has direct implications in fish growth (Hardy and Kaushik 2021; John and Mahajan 1979; Lin, Lin, and Shiau 2011; Miao et al. 2013). The dietary requirement of folic acid in fish emphasises its role in not only growth performance but also diverse functions such as immune responses (Badran and Ali 2021; Trichet 2010). Pathways involved in mitotic cell cycle and development (e.g., regulation of mitotic cell cycle, embryonic, myotome development) together with multicellular organismal water homeostasis, form a large part of the list. These pathways have in common that they relate to a biological process called “oocyte maturation”. In fish, oocyte maturation takes place before ovulation and is necessary for a successful fertilisation (Nagahama and Yamashita 2008), which may be indirectly linked to growth.

**Table 2.**
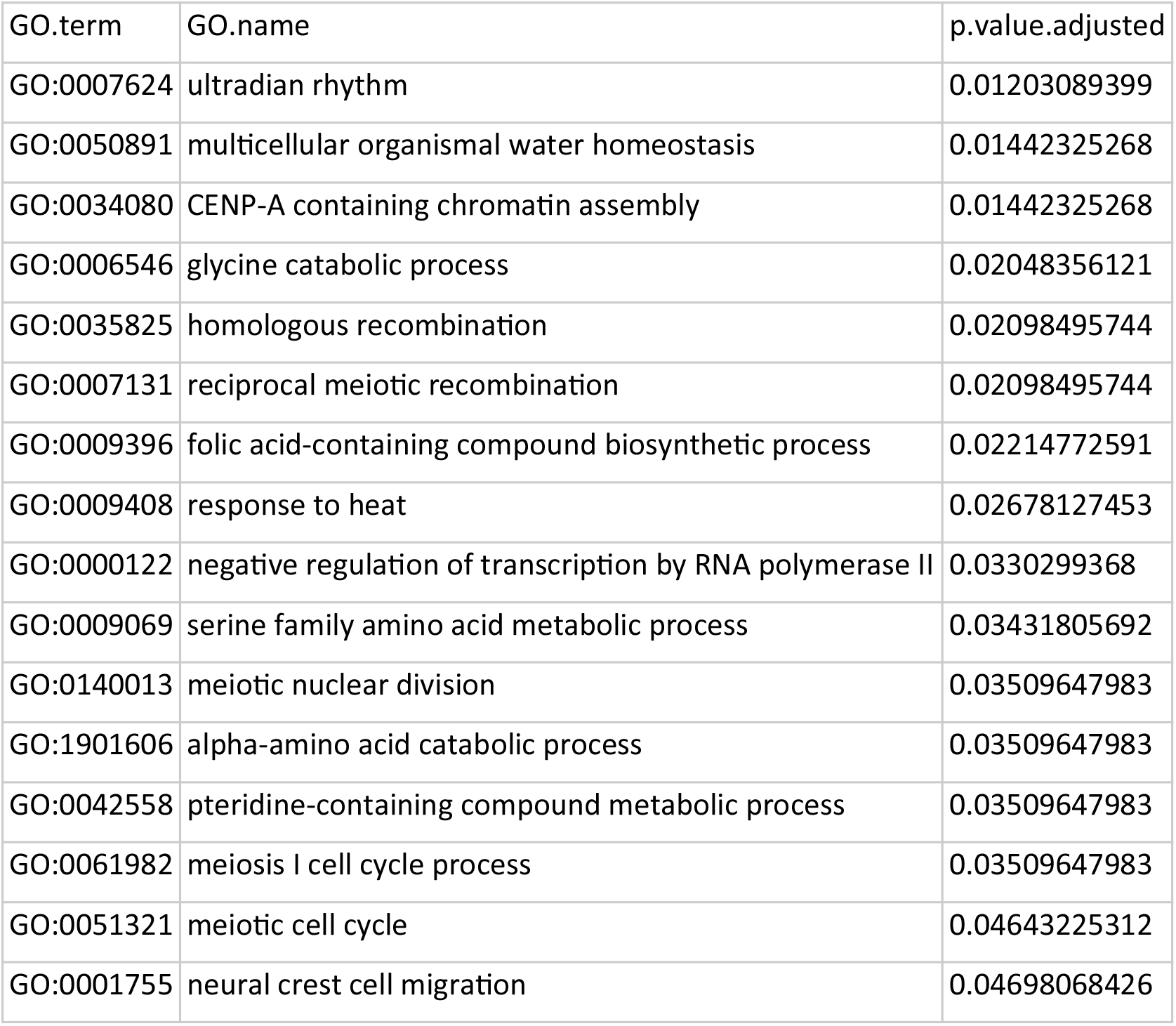
A list of enriched GO terms using genes within F_st_ windows that overlap with GWA outlier SNPs. To identify loci which are highly correlated with growth performance and selected over time, the intersections of F_st_ outlier windows and GWA outlier SNPs were investigated. When a GWA outlier SNP resides within an F_st_ outlier window, this window was counted as an overlapping outlier window (marked as red in Figure3). Any genes residing within these overlapping outlier windows were subjected to gene ontology (GO) enrichment test to identify any biological functions that correlate to growth performance and at the same time differentiated the most over time. P values are adjusted using false discovery rate. Only biological processes were presented among GO categories for the analysis.

| GO.term | GO.name | p.value.adjusted |
| --- | --- | --- |
| GO:0007624 | ultradian rhythm | 0.01203089399 |
| GO:0050891 | multicellular organismal water homeostasis | 0.01442325268 |
| GO:0034080 | CENP-A containing chromatin assembly | 0.01442325268 |
| GO:0006546 | glycine catabolic process | 0.02048356121 |
| GO:0035825 | homologous recombination | 0.02098495744 |
| GO:0007131 | reciprocal meiotic recombination | 0.02098495744 |
| GO:0009396 | folic acid-containing compound biosynthetic process | 0.02214772591 |
| GO:0009408 | response to heat | 0.02678127453 |
| GO:0000122 | negative regulation of transcription by RNA polymerase II | 0.0330299368 |
| GO:0009069 | serine family amino acid metabolic process | 0.03431805692 |
| GO:0140013 | meiotic nuclear division | 0.03509647983 |
| GO:1901606 | alpha-amino acid catabolic process | 0.03509647983 |
| GO:0042558 | pteridine-containing compound metabolic process | 0.03509647983 |
| GO:0061982 | meiosis I cell cycle process | 0.03509647983 |
| GO:0051321 | meiotic cell cycle | 0.04643225312 |
| GO:0001755 | neural crest cell migration | 0.04698068426 |

### Regions of Temporal Selection

Along with selection signals observed in regions associated with growth phenotype, we were able to identify selection signatures in other parts of the genome. In the F_st_ selection scan, pronounced high F_st_ values in LG2 and LG12 as well as low values in LG7, where previously reported inversions reside (marked in pink), were observed along with highly conspicuous deviations of **π** values (Figure S6). Thus, we calculated the frequency of a haplotype for each inversion in the temporal populations. Interestingly, only the inversion in LG12 was decreasing in its frequency consistently over time (Mann-Kendall test for monotonic trend: p-value = 0.03) (Figure 4). Within this inversion, another block of inverted region, so called “double crossover” (DC), was reported to be private to the EBC population (Matschiner et al. 2022). Thus, we identified the DC within the inversion in our sequence data (Figure S9) to examine the temporal trend of its frequency. Unlike the consistent decrease in the haplotype frequency of LG12, frequency of the DC within only decreases until 2014 then picks up in 2019. So, it seems that while the large inversion in LG12 behaves under directional selection as a whole, the DC escapes from this selection and is rather either drifting or under balancing selection on its own.

**Figure 4.**
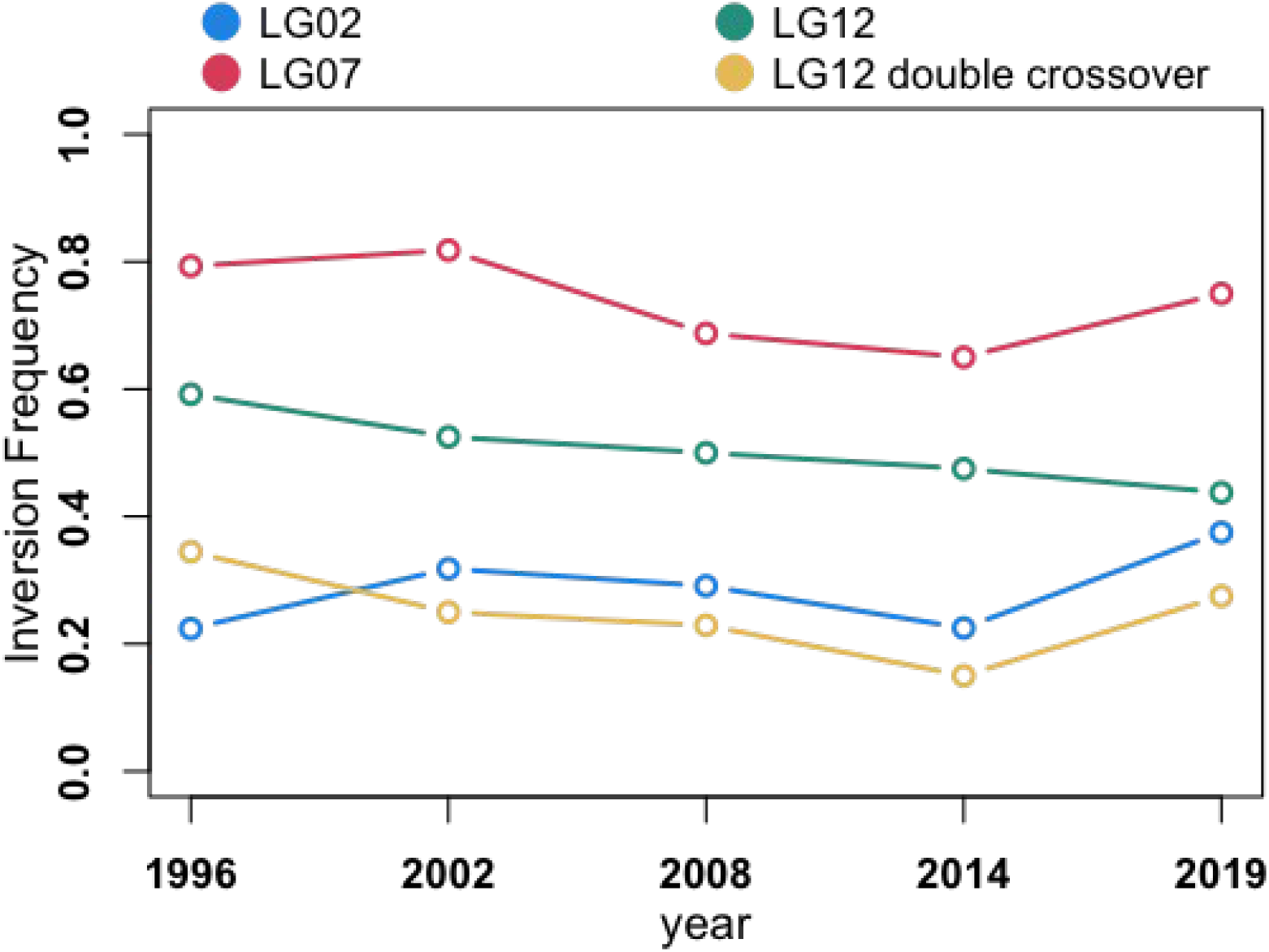
Frequency change of inversion haplotypes. The frequency of ancestral allele of each inverted region in LG2, LG7, and LG12, and the double crossover within LG12 are plotted over study period. The inversions in LG2 and LG7 display an inconsistent pattern. For the inversion in LG12, a monotonic decrease in frequency over time is observed that is statistically significant (Mann-Kendall test for monotonic trend: p-value = 0.03), whereas the frequency of double crossover within the region changes independently.

Additionally, the highest F_st_ values outside inverted regions were spread across the genome, some of which appear in peaks of clustered outliers. GO term enrichment analysis was conducted using 575 genes residing within the outlier windows of top 5% (Table S3). Several GO terms were enriched in sub-categories, which are highly related to the growth of a fish. For example, metabolisms and processing of macromolecules such as amino acids, fatty acids, and carbohydrates which in turn are key to any growth process. Fatty acid oxidation is strongly related to use of energy sources in response to feeding conditions (J. Ø. Hansen et al. 2008; Stubhaug, Lie, and Torstensen 2007; Turchini and Francis 2009) and cAMP biosynthesis is part of processing ATP, which is also critical in regulations of hormones involved in metabolism and reproductions (Miki, Van Heerden, and Fitzpatrick 1997; Takahashi and Ogiwara 2023). Also, regulation of TOR pathway, which is crucial in sensing growth hormone, nutrient or oxygen condition (Dobrenel et al. 2016; Hietakangas and Cohen 2009), was found to be enriched. As expected, some enriched biological pathways do not always show direct relevance to growth. Other highly represented clusters of GO term are in regard to developments such as gastrulation, convergent extension involved in axis elongation, and tissue morphogenesis. Interestingly, regulation of neural retina development together with melanosome transport, which is involved retinal pigmentation, may suggest temporal differentiation in the visual sensory system in EBC which is potentially relevant to depth adaptation, thus vertical movements (P. R. Berg et al. 2017; Pampoulie et al. 2015).

## DISCUSSION

This study identifies, for the first time to our knowledge in an exploited marine fish population, the genomic regions with associated gene functions that are linked to growth impairment. Reassuringly, they were also found to be under directional selection using genome scans and temporal covariance approaches. Temporal selection was likely driven by strong and documented overfishing on Eastern Baltic cod that ultimately led to the life-history change by fisheries induced evolution. A drastic decrease in the individual growth is accompanied by the contributing loci demonstrating clear evidence of directional selection with significantly positive temporal autocovariances of allele frequency changes and an excess number of overlaps with regions of high F_st_. The combination of a selection test and GWA used here is a powerful implementation of detecting an adaptive polygenic trait (Bosse et al. 2017; Brennan et al. 2018; Barghi, Hermisson, and Schlötterer 2020), which have responded to a selective pressure. In addition, observed directional change in the frequency of ancestral haplotype of inversion in LG12 but not for its double crossover region underscores the heterogeneous response of the genome under selection.

### Overall Patterns of Temporal Genomic Change

Non-significant change of nucleotide diversity, a lack of clustering pattern in PCA, and genome-wide covariance patterns resembling neutral population suggest that migration, gene flow and other non-adaptive processes were negligible over the study period, at least not at the resolution provided by the methods employed. In addition, heterogeneous response of the genome, by utilising standing genetic variation across the genome, may have been driving the changes in phenotype potentially through different metabolic processes (Crespel et al. 2021). Hence, the premise of this study, namely that EBC is a closed, self-sustained gene pool without immigration of divergent genotypes, is supported. Moreover, the possibility of other traits undergoing selection or drift in divergent directions than the targeted trait, could potentially obscure the genome-wide signal of size selective fishing in wild populations. For example, in EBC, an opposing selection pressure against small female body size can be hypothesised. This is because larger females produce larger and more buoyant eggs that permit them to float higher in the water column (Nissling and Vallin 1996), away from the near-bottom where the oxygen conditions are worsening.

Absence of evidence of overall pattern does not equate to evidence of absence of selection (Bosse et al. 2017; Fuller et al. 2020). Despite the lack of overall pattern, evident non-random signals were observed when targeting specific regions, the inverted region of LG12 and the candidate loci of GWA. Against the background of no overall change in genomic patterns (Figure S5 and S6) the directional change in the frequency of inversion in LG12 clearly suggests selection in parallel to the apparent decline of growth rates. In EBC, apart from some adaptive loci linked to salinity and oxygen found within the chromosomal inversion in LG2 (Paul R. Berg et al. 2015), any evidence on adaptive or ecological roles of inversion haplotypes is generally lacking. Although no GO term was found significantly enriched for genes located within the inverted region of LG12, the ancestral homozygous status of individuals, together with body size, had a correlation to lower survival rate in an Atlantic cod population in the North Sea (Barth et al. 2019). In addition, SNP loci within this inverted region were highly correlated with temperature and oxygen level at the surface likely driving the differentiation of cod populations (Paul R. Berg et al. 2015). Interestingly, the frequency of double crossover (DC) within the inverted region seems to be fluctuating independently of the large inversion. This region is densely packed with genes including three vitellogenin genes, which are crucial for creating buoyancy of eggs for the survival and successful spawning in EBC (Nissling and Westin 1991). In this context, we speculate that the selection pressure acts upon the inversion as a whole, but is relaxed for the crucial set of genes in the DC by broken linkage disequilibrium. This scenario might also explain the hypothesis of the opposite selection pressure on body size of females mentioned above.

### Functional relevance of selected loci

Several enriched GO pathways for the overlapping regions of GWA and F_st_ outliers suggest that the selected gene functions are causally linked to altered growth in EBC (Table 1). Light manipulation to tweak the ultradian rhythm of individuals, thus the long term seasonality, is a very common method to control growth and maturity in fish aquaculture including Atlantic cod (Skulstad et al. 2013; Taranger et al. 2010; Karlsen et al. 2006; T. Hansen et al. 2001). Depending on the applied photoperiod, sexual maturation can be controlled, either postponed or advanced, which is tightly entangled to somatic growth of a fish (T. Hansen et al. 2001; Davie, Porter, and Bromage 2003). In addition, water homeostasis is important in egg hydration during the oocyte maturation process to make floaty eggs, which is one of the major evolutionary acquisitions for pelagic teleost fish (Fyhn et al. 1999). Oocyte maturation takes place before ovulation and is necessary for a successful fertilisation (Nagahama and Yamashita 2008). Specific hypotheses directly connecting oocyte maturation and growth are currently lacking in the field. Nevertheless, it is well conceivable that the timing of spawning, through control of oocyte maturation, may be critical for successful reproduction, as maturation process is highly affected by energy allocation (Roff 1993), thus tightly linked to somatic growth in a fish’s lifespan. Lastly, the biological process of “response to heat” is indeed highly linked to growth traits in fish. Warmer temperatures as the Baltic sea has been experiencing (Meier et al. 2022), critically impact the species throughout the lifespan from larva to adult stage (Oomen et al. 2022; Righton et al. 2010) and are dynamically interlinked with other environmental factors such as oxygen. Thus, it may suggest that the slow growth trait was also mediated or accompanied by shifts in temperature response over time.

In spite of the obvious, functionally aligned links to growth from candidate loci, there seems to be a general lack of congruency in the genetic contents compared to previous studies which experimentally addressed the genomic effects of size-selective harvest selection. Therkildsen et al. (2019) resequenced samples from the seminal study of Conover and Munch (2002) that subjected Atlantic silversides to 5 generations of upwards and downward selection with respect to body size. They listed enriched GO terms from highly differentiated loci accompanied by body size changes under different harvest regimes. Data from the present study found no intersection to above results, which is perhaps not surprising given that Therkildsen et al. (2019) itself observed highly divergent genomic responses across replicates under the same treatment. In another experimental study in zebrafish, Uusi-Heikkila (2015) identified another set of genes selected by fishing pressure that were also not present among the genes listed as outliers in this study. Lastly, the *vgll3* and *six6* genes that are of high effective size in age at maturity in salmonids species (Barson et al. 2015; Ayllon et al. 2015), a tightly linked yet different life history trait, were not found to be significant in any of the present analyses. This lack of consistent patterns of identified genes and pathways in this study compared to previous studies of FIE as well as among the studies indicate that there are heterogeneous responses in the genome level either under same phenotype changes, growth, or under same selective pressure, size-selective fishing.

### Future directions and Implications in Fisheries Management

With promising results showcasing EBC as an evolving population with stunted growth, this study directs to important future research agendas and implications in managing the stock. First, it is important to note that these evolutionary responses occurred in the context of dynamic interplays of fisheries and adverse environmental factors. Examples abound that overexploitation will cause evolutionary change, but these responses are always highly context dependent. Depending on the life history traits under selection and their genomic architecture, strength, length, and types of selection pressure together with natural selection by various environmental factors may be reinforcing or counteracting the trait evolution in a convoluting manner. Environmental factors, such as hypoxia and temperature increase as well as ecological variables, such as prey and predator interactions and inter- species competition in the Baltic Sea, have been directly and indirectly influencing the population at the same time (Casini et al. 2016; Eero et al. 2012; Limburg and Casini 2018; 2019; Neuenfeldt et al. 2020), which may or may not have exerted an evolutionary pressure. For example, sea surface temperature has risen around 1.5 °C during the study period (Siegel and Gerth, 2018) and can only maximally explain a 6 % decrease in body size according to gill-oxygen limitation theory (Pauly and Cheung 2018). Then the hypoxia in the bottom water in the Baltic Sea has been continuously deteriorating since the 1930s and the extent to which Bornholm Basin has been directly impacted were variable depending on inflow from North Sea and surrounding rivers (Carstensen et al. 2014; Stockmayer and Lehmann 2023). Thus, given time-series data of an adequate resolution, direct associations of genotypes and fishing pressure as well as environmental variables are an essential further step to take.

Secondly, this study focuses on the critical period of a steep decline and lowest point in growth from 1996 to 2019 and provides a contemporary snapshot of the long-term population dynamics of EBC. However, it is crucial to acknowledge that growth has fluctuated, with an increase during the 1960s to 1980s, followed by a noticeable decline from the 1990s to the present (Mion et al. 2021). Thus, the direct causes and evolutionary responses shaping the growth trend warrant further investigation within a longer timeframe, preceding and succeeding the study period. Especially, when an inherent lag in evolutionary response, referred to as “Darwinian debt” (Ulf Dieckman in an interview by Cookson, *Financial Times*) may be contributing to the delay of recovery by compromising growth potentials and population resilience (Anderson et al. 2008; Ahti, Kuparinen, and Uusi-Heikkilä 2020), it urges a comprehensive examination of long-term ecological and evolutionary consequences.

Lastly, successful management plans for EBC must incorporate evolutionary aspects into their framework, e.g. introducing F_evol_ (Hutchings 2009), integrating evolutionary processes into economic assessments of management plans (Eikeset et al. 2013; Schenk, Zimmermann, and Quaas 2023). Having said that, the impact of such measures on fisheries management may be limited at this stage as the damage has already been done. At present, the evolutionary debt has been accumulated and despite the current moratorium, the stock recovery falls short of expectations due to concurrent contribution of ecological and environmental factors to stock condition (Eero et al. 2023). Whether this lack of recovery is already one consequence of the Darwinian debt is an interesting hypothesis to explore in the future.

## MATERIALS AND METHODS

### Sample Collections

In total 152 cod individuals were used in this study after excluding individuals which were identified as either a western Baltic cod from genetic analysis (9 samples), an outlier from growth analysis with measurement errors (1 samples), and of low sequencing quality (2 samples). Sampling was done in two different ways to cover the available time period and the full range of phenotype in the sampling pool. 1) A set of samples, called “random” hereafter, were randomly sampled along the length distribution for five catch years; 31 from 1996, 22 from 2002, 24 from 2008, 20 from 2014, and 20 from 2019. 2) As another set of samples, called “phenotype” hereafter, 19 smallest mature fish and 18 largest immature fish were selected from the catch year 1996-1998. As any age information of the archived samples was not available, neither sample based on the cohort nor on length at first maturity was possible. The rationale was that by sampling immature fish, which would be first mature in the following year if they had not been caught, and small, presumably young, mature fish, we attempted to cover as wide a range of phenotype variation as possible.

Otoliths and finclips were collected in the Baltic Sea Integrative Long-Term Data Series of the research division Marine Evolutionary Ecology at GEOMAR, carried out annually since 1996. They were taken on board from cod caught in Bornholm Basin (Figure 1A), of which their phenotype data (e.g., body length, weight, maturity stage, and sex) was recorded (Table S1). Otoliths were stored in paper bags. Finclips were stored in 100% ethanol at -20 °.

### Age Reading of Otoliths

As the conventional otolith reading method has not been reliable for EBC, a newly developed method was employed to acquire age information of the sequenced samples in order to model growth based on Hüssy et al. 2021. For chemical analysis, otoliths were embedded in Epoxy resin (Struers®) and cut to have exposed surface of the core and the rostral part. Trace element analysis were conducted by Laser Ablation Inductively Coupled Plasma Mass Spectrometry (LA-ICP-MS) to measure magnesium (^25^Mg), phosphorus (^31^P), and calcium (^43^Ca), which exhibit seasonal variations in EBC (Heimbrand et al. 2020; Hüssy et al. 2021). Since the elements were read from the core of an otolith to the edge, the measured element traces represent the chemical characteristics of an individual’s lifespan from the hatch to catch. With the measured element profile, a statistical analysis was carried out to determine the age. Chemical minima were identified using local polynomial regression function “loess” and “peaks” in R (R Development Core Team, 2022). The arguments were set based on the settings used in age reading of tag-and-recapture cod samples in previous studies. The numbers of minima in Mg and P, which suggest the fish’s exposure to the coldest temperature of a year (February and March), are counted as the age of an individual (Figure 2 in Hüssy et al. 2021). When the two values disagreed, the element profiles were visually examined. This approach is not as stable for the signals near the otolith edge. Thus, visual assessment was conducted for the samples caught in the first quarter of a year. As a result, annual chemical radii for each individual, total otolith radius, as well as the age at catch were extracted. The exact details of preparation of otoliths, procedures concerning LA-ICP-MS, and the statistical analysis can be found in Hüssy et al. 2021.

### Modelling Individual Growth Rates

To acquire a heritable phenotype that may have been affected by fishing pressure, we modelled individual growth using the age information. Although it was recently confirmed that the growth of EBC has impaired over last decades (Mion et al. 2021), it is crucial to obtain the growth pattern of sequenced individuals to integrate genotypes and phenotype.

To fully utilise the hierarchical nature of the estimated otolith chemical annuli at age of fish individuals from different catch years, Bayesian hierarchical modelling was applied using R2jags v0.7.1 R package (Su and Yajima 2021). The von Bertalanffy growth function (von Bertalanffy 1957) was fitted to distance from core to chemical annuli at age on otoliths:

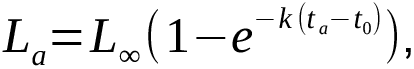

where *L_a_* is distance from otolith core to each chemical annulus, *t_a_* is the estimated age at the annulus, *L∞* is asymptotic length in an otolith scale, which is hypothetical otolith length at age of infinity, *k* is a growth coefficient, and *t_0_* is hypothetical age when length equals zero. Three levels of hierarchy included measurements of annuli at age, nested in a fish individual, again nested in a group of a catch year. As a result, *L∞* and *k* parameters were estimated for each individual and also each catch year. We took the most conservative approach of priors, applying a gamma distribution for catch years and normal distribution for individuals with relaxed standard deviations (details in the script). To fit the model, 100,000 iterations were observed for three MCMC chains and the first 10,000 were discarded as burn-in. The median of Rhat values were 1.0036 and model convergence of the chains were visually examined in addition (Figure S10). As an additional assessment of the model, residuals were calculated from estimated otolith length from the model and observed length of otolith annuli (Figure S11). Here, the variance of residuals is larger for the first year which could be caused by the uneven number of observations that were fed to the model for each age. Nevertheless, the overall residuals remain near zero for all years. To avoid any bias of condition towards bigger fish, relative condition factor (Le Cren 1951) was used to test whether fish condition could predict any of the growth parameters and Φ. Back-calculation of fish length was conducted using an equation from (Hüssy, Eero, and Radtke 2018), using biological intercepts specific (Campana 1990) for Baltic cod. Accordingly, *L_0_*, which is the fish length at age 0, was set to 4.3 and *O_0_*, the otolith length at age 0 was set to 0.01.

### DNA Extractions

For genetic materials, DNA was extracted using otoliths from earlier years (1996- 1998, 2002, and 2008) and fin clips from recent years, 2014 and 2019. Otoliths and finclips were always handled with tools (e.g., forceps) which were cleaned with ethanol 70% and sterilised in between each individual sample to avoid cross contamination. The extraction procedure for both otoliths and finclips were conducted following the standard protocols from either DNeasy® Blood & Tissue Kit (Qiagen, Aarhus, Denmark) or NucleoSpin® Tissue Kit (Macherey-Nagel, Düren, Germany). Otoliths were fully submerged in the lysis buffer to lysate any remnant tissues then removed from the buffer. The lysate then was treated as in the manuals provided by the kits. Fin clips were cut into small pieces (up to 25mg), submerged in a lysis buffer, then continued following the protocols. The extracted DNA was purified using Qiagen QIAquick® PCR Purification Kit (Qiagen, Aarhus, Denmark). DNA quality was checked with standard electrophoresis in 1% agarose gel and the quantity was measured using NanoDrop^TM^ and Qubit Assay (Thermo Fisher Scientific^TM^, Carlsbad, USA).

To validate cross contamination that might have occurred during the sample collection, archiving process, and DNA extraction, microsatellite (MSAT) analysis was done for DNA extracted from otolith samples. Four MSAT sites were used. A multiplex PCR was conducted with four primer pairs on a 96-well plate. The PCR product was mixed with Hi-Di^TM^ mix (Thermo Fisher Applied Biosystems^TM^, Carlsbad, USA) with GeneScan^tm^ LIZ dye Size standard^tm^ (Thermo Fisher Applied Biosystems^TM^, Carlsbad, USA). Capillary electrophoresis was done with the reaction mix using ABI PRISM 3100 Genetic Analyzer (Thermo Fisher Applied Biosystems^TM^, Carlsbad, USA). The MSAT peaks were analysed using GeneMarker® software (Softgenetics, State College, USA). As the chosen MSAT loci typically show more than ten alleles per site in a population, when samples are mixed the likelihoods of encompassing the same allele at a single MSAT locus are small and are virtually zero if several such loci are combined. Thus, samples showing multiple peaks for any MSAT locus were identified as cross contaminated and subsequently excluded from the data set (see examples in Figure S12).

### Library Preparation and Sequencing

2x100 bp paired end library preparation for 16 samples from 1996 was done in the Ancient DNA Laboratory at the Institute of Clinical Molecular Biology (IKMB) as a pilot to check if they should be treated specially like historic DNA samples. The details of the manual library preparation can be found in the method section in Krause-Kyora et al. 2018. For the finclip samples from 2014 and 2019, 2x150bp paired end libraries were prepared using Illumina DNA Prep kit (Illumina, San Diego, USA) by the Competence Centre for Genomic Analysis (CCGA) Kiel. These libraries (16 otolith samples from 1996 from pilot and 40 finclip samples from 2014 and 2019) were sequenced on Illumina 6000 S4 Flowcell (Illumina, San Diego, USA) by CCGA Kiel. In the end it was concluded that older otolith samples can be treated the same as the rest, yielding sequence data of comparable quality. Thus, rest of the samples, including “phenotype” samples from 1996-1998 and “random” samples of 1996, 2002 and 2008, were sent to Norwegian sequencing center (NSC) for 2x150 bp library preparation using Illumina Nextera DNA library preparation kit (Illumina, San Diego, USA) followed by sequencing on Illumina NovaSeq S4 Flowcell (Illumina, San Diego, USA).

### Read Processing and Variant Calling

All sequenced reads from this study were processed together with published population data from Barth et al. 2019, to include 23 EBC (named BOR), 22 WBC (KIE), and 24 North Sea (NOR) cod samples, which were later partitioned out. This was to identify WBC in our samples and test for any sequencing bias in our samples (Figure S13) as well as to conduct ancestry painting, which includes WBC and EBC individuals of known inversion status as reference (explained below). All sequenced reads were processed following the GATK best Practices workflow by Broad Institute (GATK v4.1.9.0) (Van Der Auwera et al. 2013). All the detailed commands, parameters, and filtering options in the bioinformatics workflow are included in the provided git repository. Mapping to the reference genome of Atlantic cod, gadMor3.0 (NCBI accession ID: GCF_902167405.1), the median coverage of each individual ranged from 4x to 31x with a median of 12x for all samples. Two samples from 1996 were excluded based on their low mapping coverage below 4x.

After variant calling, raw SNP variants were first hard filtered based on different qualities of variant sites according to best practices. Then, only biallelic SNPs were selected and filtered again based on genotyping quality, missingness, read depths, and minor allele frequency (MAF) of 0.005 to produce the final variant call file in a vcf format containing 5,847,389 variants. When possible, this full set of variants based on MAF > 0.005 were used, although some analyses were carried out using 4,685,343 variants filtered with MAF>0.01 due to the processing time and resource limitation.

Further analyses were done with two separate sets of variants resulting from different partitioning of the total sample set (also partitioned from WBC and North Sea samples), as parts of the sampling was intentionally biased for “phenotype” samples as explained earlier. i) 115 of “random” samples were used for the analysis identifying signatures of selection over time. ii) A total of 152 samples including “random” and “phenotype” samples were used for genotype-phenotype association. The subset of the master vcf file was created using bcftools v1.2 (Danecek et al. 2021) then fixed sites were removed using GATK SelectVariants (v4.1.9.0).

### Population Statistics and Principal Component Analysis

To examine any temporal differentiation in EBC independent of phenotypic data, 115 “random” samples were used to compute Nucleotide diversity ( *π*), between population nucleotide divergence (dxy), F_st_ and principal component analysis (PCA). For calculating *π* and dxy, guides provided by Pixy (1.2.7.beta1) (Korunes and Samuk 2021) were followed. A vcf file containing invariant sites was created, using GATK GenotypeGVCFs with option –all- sites followed by site filtering steps using GATK VariantFiltration with same criteria as in hard filtering of variants and followed by vcftools v0.1.16 (Danecek et al. 2011) on missingness of 0.8 and mean read depths of 10. This filtered all-site file was combined with the final variant file to create the input vcf file for Pixy. A total of 81,462,138 records including invariant and variant sites, were used to calculate *π* for each catch year and pairwise dxy in 50kb non-overlapping windows. For genome-wide nucleotide diversity for each temporal population, average *π* value for all windows was calculated according to the equation provided by Pixy.

PCA on the subset of SNPs (4,685,343 after filtering for MAF > 0.01) was carried out using the R package pcadapt v4.3.3 (Privé et al. 2020). Scree plots of total variance explained by each principal component (PC) were examined to decide up to which PCs to investigate. When all sites were included, a unique clustering pattern driven by inversion status of individuals appeared (Figure S14). Thus, sites within the inverted regions (identified as described in *Identifying inversion status)* were excluded then pruned based on linkage disequilibrium (2,030,929 SNPs) to examine the remaining population structure.

Weir and Cockerham’s F_st_ was calculated using vcftools v0.1.16 in 20kb windows. Only weighted F_st_ was used for plotting and interpretation of the data. All plots were created in R (R Development Core Team, 2022) using the base “plot” function.

### Genome-wide Temporal Covariance and Simulation

Genome-wide temporal covariance was calculated using a modified python script in Jupyter notebook based on the functions in cvtkpy (http://github.com/vsbuffalo/cvtk) published in Buffalo and Coop (2020). Error bars were calculated by bootstrapping covariance values, resampling blocks of loci 5000 times, using the bootstrap function provided by cvtkpy. As initial genome-wide temporal covariance showed an inconclusive pattern, we simulated a neutrally evolving population to compare the covariance values as a control. First, backward-in-time simulation was employed to create a population with matching diversity using msprime v1.2 (Baumdicker et al. 2022), with mutation rate 3.5e-9, recombination rate 3.11e-8, 5000 genomes, and a sequence length of 30Mb. With this population as a founding population, a forward-in-time simulation was conducted using SLiM v2 (Haller and Messer 2017). Additional 100 generations were burned in at the beginning of the simulated time. From generation 101, 20 individuals were sampled from the simulated population for five generations, to imitate the sampling scheme of wild population. Final vcf file was created to calculate the covariance of the simulated temporal populations. This was replicated 100 times to create a distribution of patterns from neutrally evolving populations.

For the calculation of temporal covariance, a custom script in R language was used which replicated the functions in cvtkpy.

### Genome-wide association analysis (GWA)

To identify specific genomic regions responsible for growth variation in the EBC population, genome-wide association study was conducted. Growth performance was converted into an index using the growth estimates, Φ = log *k* +2 log *L∞* (Moreau, Bambino, and Pauly 1986; Munro and Pauly 1983). Subsequently, this variable was subjected to a univariate nonlinear mixed model to identify loci associated with the growth change using GEMMA v0.98.3 (Zhou and Stephens 2012). A total of 679,584 SNPs were used after filtering for minor allele frequency of 0.05 and missingness of 0.1 as recommended by the developers. Genetic population structure was considered as a random effect and sex as covariates to incorporate and eliminate possible other contributing factors. Genomic inflation factors and QQ plots showed that systematic biases were adequately corrected from the other contributing factors (Figure S15). After correcting for multiple testing, using false discovery rate (Benjamini and Hochberg 1995), with the number SNPs sites not in linkage disequilibrium (174,541), there were no SNP sites with genome-wide significance for Wald test p-values observed. Instead, as an exploratory approach to identify the loci that are most likely to be associated with growth, a cutoff which includes the most obvious peaks but excludes more spurious signals in the Manhattan plot were set. As results, SNP loci occupying the 0.05% tail of distribution of the p-values, 338 variants, were assigned as outliers for further analysis (referred to as “GWA outliers”).

### Calculating and Bootstrapping Temporal Autocovariance of GWA outliers

To demonstrate the directional changes over time in allele frequencies of the GWA outliers which are accountable for the growth variations, temporal covariance of the outlier loci was calculated in R. We used delta values of different time windows, lag-2 and lag-3, contrary to those with lag-1 provided in the *cvtkpy* package, which always uses consecutive time points to calculate the allele frequency changes. This was to avoid including a shared time point in calculating autocovariance which showed positive covariance values in the simulated neutral populations and was likely driven by the shared time point rather than a true signal of selection. To assess the significance of observed covariance, a permutation test was conducted calculating temporal covariance values using 338 random loci sampled from all SNP sites in GWA analysis. The observed values were compared to the distribution of 1000 random permutations.

### Gene Identification and Gene Ontology (GO) Term Analysis

To further assess the biological relevance of any outlier loci or windows from genomic analysis, two approaches were employed, 1) by searching for functional annotations in targeted genes for GWA outlier SNPs and 2) by gene ontology (GO) term enrichment analysis using a set of outliers. For 1), among the 338 SNPs assigned as GWA outliers, only regions with clustering outliers with flanking SNPs with low values (marked with red arrows in Figure 12) were examined in depth. Genes located at or within 5 Kb up- and downstream of the outliers were further searched for their biological functions in the literature. The search was carried out using the gene names or descriptions, targeted with or without key words, e.g., fish, growth, maturity, and reproduction to find the most relevant functions to this study. Genes were listed by cross referencing each SNP to annotated genes in the gadMor3.0 annotation database ("gmorhua_gene_ensembl") in Ensembl using the BioMart v2.54.1 R package (Durinck et al. 2005). Same database and workflow were used in identifying genes lying within F_st_ outlier windows and in overlapping windows of F_st_ and GWA outliers. With the listed sets of genes, enriched GO terms were identified using the GO terms provided in the annotations of the gadMor3.0 database as “universe.” The workflow was based on the vignette provided by GOstats v2.64.0 R package (Falcon and Gentleman 2007).

### Identifying Inversion Status

Four large (5-17 Mbp) chromosomal inversions in Atlantic cod species have been previously identified (Paul R. Berg et al. 2015; Kirubakaran et al. 2016; Sodeland et al. 2016), three of which are polymorphic in the EBC population. We targeted these regions as candidate supergenes which may have undergone selection over the study period and examined how their frequency changed over time. With prior knowledge of inversions located in LG2, 7 and 12, PCA was done on subset vcf files of each chromosome. Three distinct clusters of individuals of different inversion status (homozygous ancestral, homozygous derived, and heterozygous, “ancestral” status adopted from Matschiner et al. 2022) were observed, which was used for individual assignment. Then, F_st_ values were calculated among these three groups (each pairwise and global) and plotted to identify boundaries of the inversions (Figure S16). These boundaries were used to subset the bedfiles to feed as input of local PCA analysis. The inversion status of individuals was verified again by visually examining local PCA plots for each inversion status (Figure S17). When ambiguous, the individuals were visually examined for their genotypes in IGV v2.12.0 (Thorvaldsdóttir, Robinson, and Mesirov 2013).

To identify the individual status of double crossover, ancestry painting was carried out following a tutorial from a git repository of M. Matschiner (github.com/mmatschiner/tutorials/tree/master/analysis_of_introgression_with_snp_data). We used four samples (homozygotes ancestral: KIE1203003, BOR1205002 and homozygotes derived: KIE1202006, KIE1203020 from Barth et al. 2019) as reference of ancestral and derived homozygotes and two EBC (BOR1205003, BOR1205007; identified in Matschiner et al. 2021) as “control” of double crossover. SNP sites between positions 6.5Mb and 7.5Mb in LG12, (Note that the location is different than reported in Matschiner et al. as different reference genomes were used) which are fixed 80% in these reference individuals, allowing for 20% of missingness, were painted two different colours in EBC individuals (Figure S9). Double crossover status, either ancestral/derived homozygous or heterozygous, was assigned by visual examination.

## ACKNOWLEDGEMENTS

## Author contributions

Conceptualization: TBHR, KYH, JD Sample Acquisition: KYH, TBHR, JD

Methodology: KYH, EEK, JD, SJ, CH, JF, BKK, KH, TBT, BDH

Investigation: KYH, RB, CM Visualization: KYH Supervision: TBHR

Writing—original draft: KYH, SJ, TBHR Writing—review & editing: all authors

## Data and materials availability

The sequence data is archived under NCBI BioProject PRJNA1128530

The scripts and metadata used in this study are archived in a Gitlab repository https://github.com/kwiyounghan/FIE_Baltic_cod

## Supporting information

All supplementary figures and tables

