## Supplementary material for "Genomic Evidence of Fisheries Induced Evolution in Eastern Baltic cod": All supplementary figures and tables

**Figure S1. Boxplots of estimated individual von Bertalanffy growth parameters over time.**

The individual level parameters were estimated in the growth model using “random” and “phenotype” samples. The posterior predictions of estimated parameters of individual level were grouped in temporal populations to calculate the median and the lower and higher whiskers. **A.** $L_\infty$, and **B.** growth coefficient k, and **C.** otolith radii at age 1 of all fish. Colour codes are based on individuals’ catch years as in the legend.

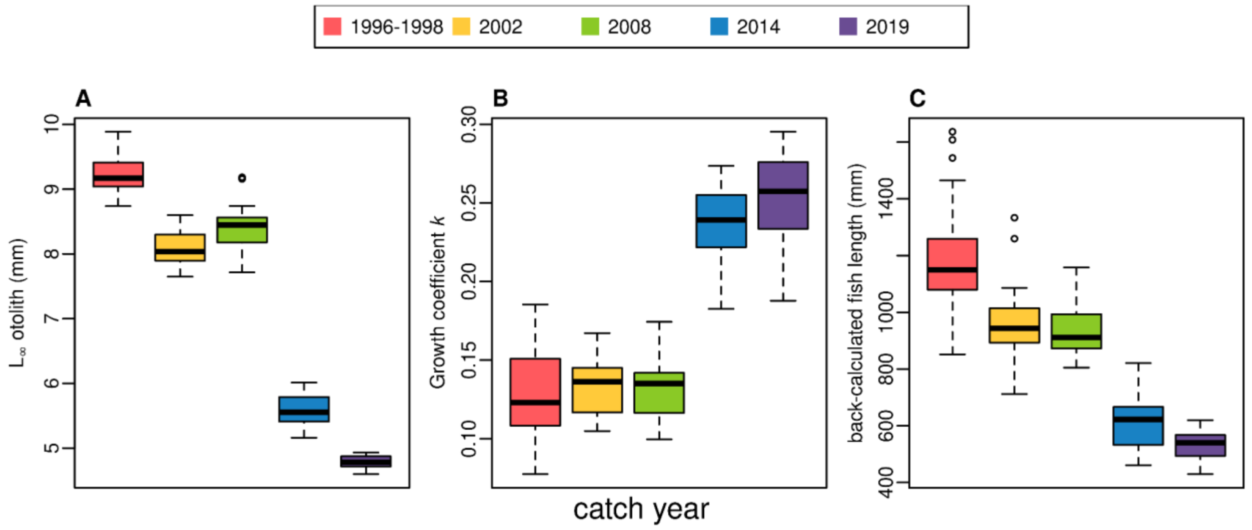

**Figure S2. Fish condition at catch and relationship to growth.**

Relative condition factor was calculated for individual using fish body weight and length at catch to test for a potential predictor for individual growth. A. Boxplot of condition for individual fish grouped based on the catch year, the temporal populations. Individual condition is plotted against B. growth coefficient *k,* C. asymptotic otolith length $L_\infty$, and D. growth performance index, Φ. The correlation coefficients for each parameter are, r = -0.03 (p > 0.05), r = 0.09 (p > 0.05), and r = 0.09 (p > 0.05).

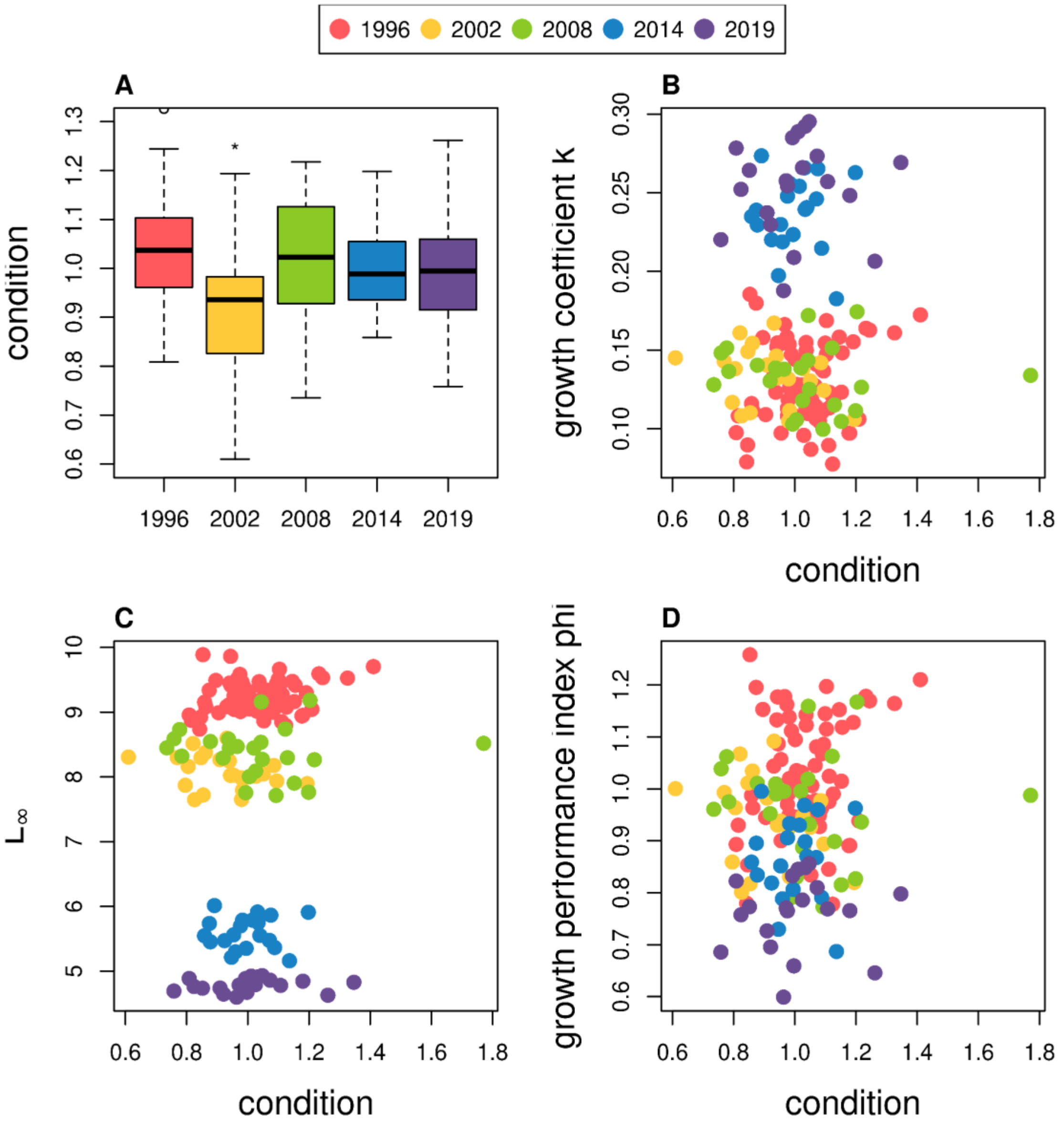

**Figure S3. Genome-wide temporal covariance analysis on SNP polymorphism through time.**

Each line in the covariance plot represents the temporal covariance, cov($\Delta p_s$, $\Delta p_t$), calculated using allele frequency changes in two time windows of *s* and *t*; the lines are coloured accordingly to rows of the temporal covariance matrix on the right side. The x-axis of the plot shows the later time windows in the calculation, $\Delta p_t$. For example, the three green points are, from left, cov(Δ1996-2002, Δ2002-2008), cov(Δ1996-2002, Δ2008-2014), and cov(Δ1996-2002, Δ2014-2019). Error bars of 95% confidence interval calculated by bootstrapping covariance are drawn as lines over each point.

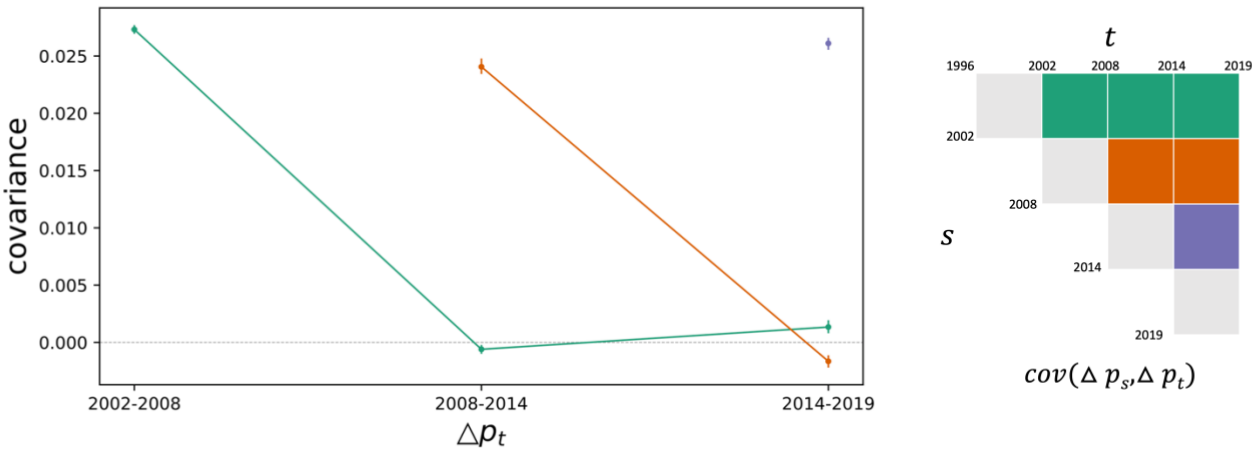

**Figure S4. Simulated genome-wide temporal covariance through time in the absence of selection.**

Similar to Figure S1, each line in the covariance plot represents the temporal covariance, cov($\Delta p_s$, $\Delta p_t$), calculated using allele frequency changes in two time windows of *s* and *t*; the lines are coloured accordingly to rows of the temporal covariance matrix on the right side. The x-axis of the plot shows the later time windows in the calculation, $\Delta p_t$. Instead of the catch years, simulated generations (t1 - t5) were used. The median of all 100 simulated populations are plotted in colours and all temporal covariance values are plotted in grey lines to show random distribution of covariance values in a scenario of neutral evolution.

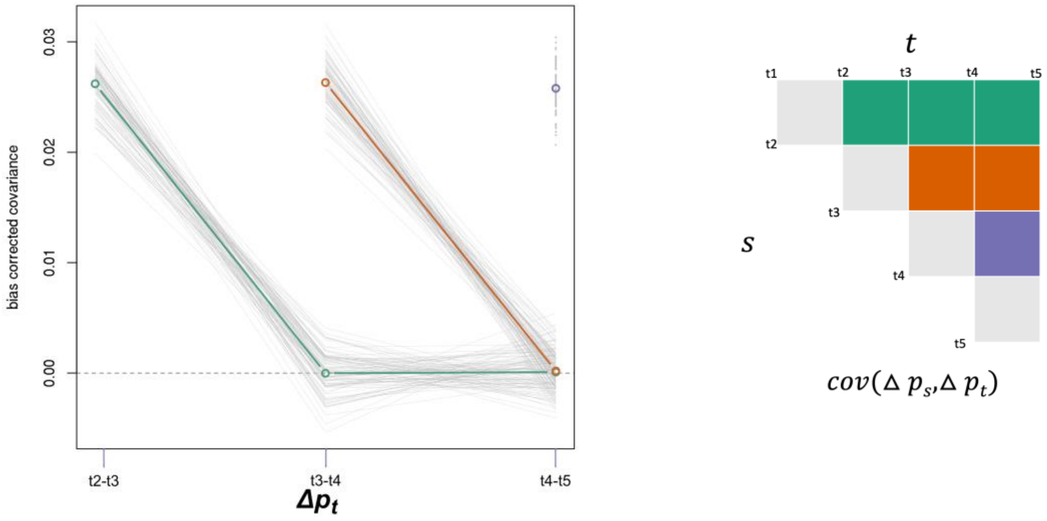

**Figure S5. Distributions of the simulated temporal covariance values compared to covariance from real data in Figure S3.**

To understand the significance of the genome-wide temporal covariance values, the values from real data (Figure S3) were compared to the simulated values (Figure S4). The histograms are frequency distribution of simulated values (100 replications) for each pairwise covariance of different time windows. The colour codes follow the covariance matrix in Figure S4 of simulated time points (T1-T5), which corresponds to five time points 1996-2019. The red lines represent the covariance values of the real allele frequency changes for corresponding time windows. For example, for the first histogram of cov(ΔT2-T1, ΔT3-T2) were compared to cov(Δ2002-1996, Δ2008-2002).

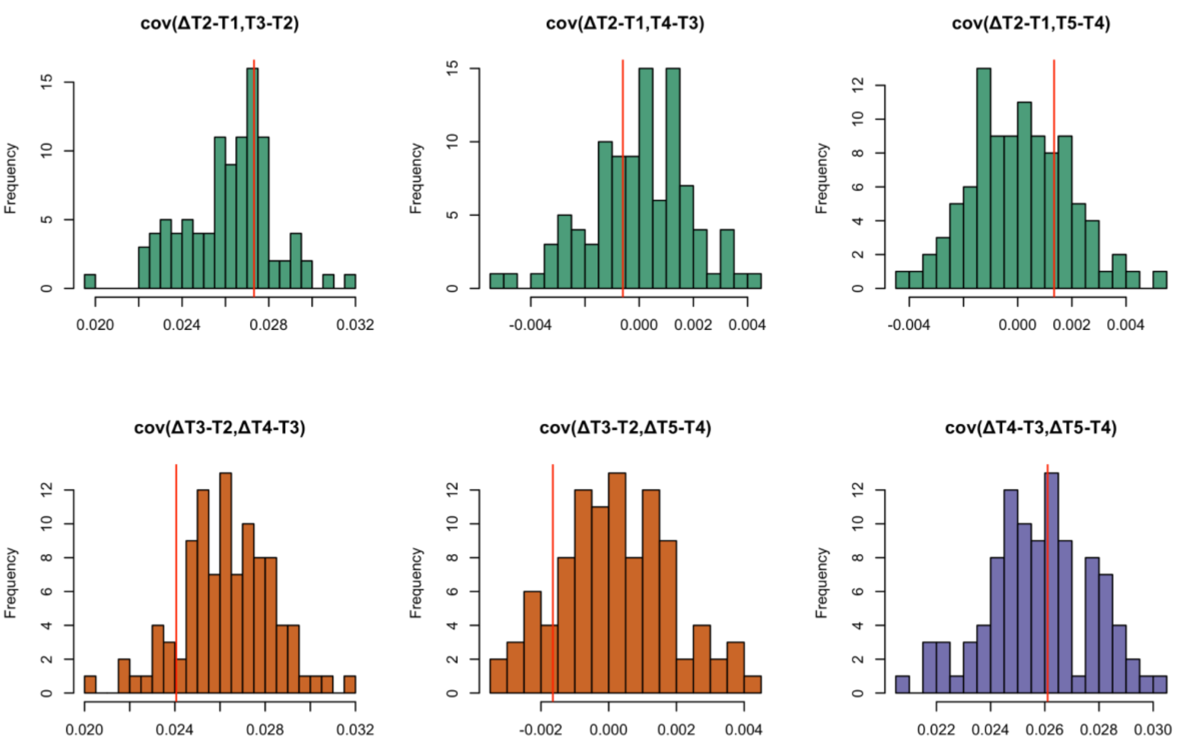

**Figure S6. Population genetic statistics in 50Kb non-overlapping windows along the genome**

**A.** Nucleotide diversity ($\pi$) for each catch year and **B** absolute divergence between populations ($d_{xy}$) of first time point 1996 to all other years. Colours as in the legends above the plots. A total of 81,462,138 invariant sites and variant SNPs were used to calculate the values. For plotting, an R function “loess.smooth” was used to smooth the curves for visual ease.

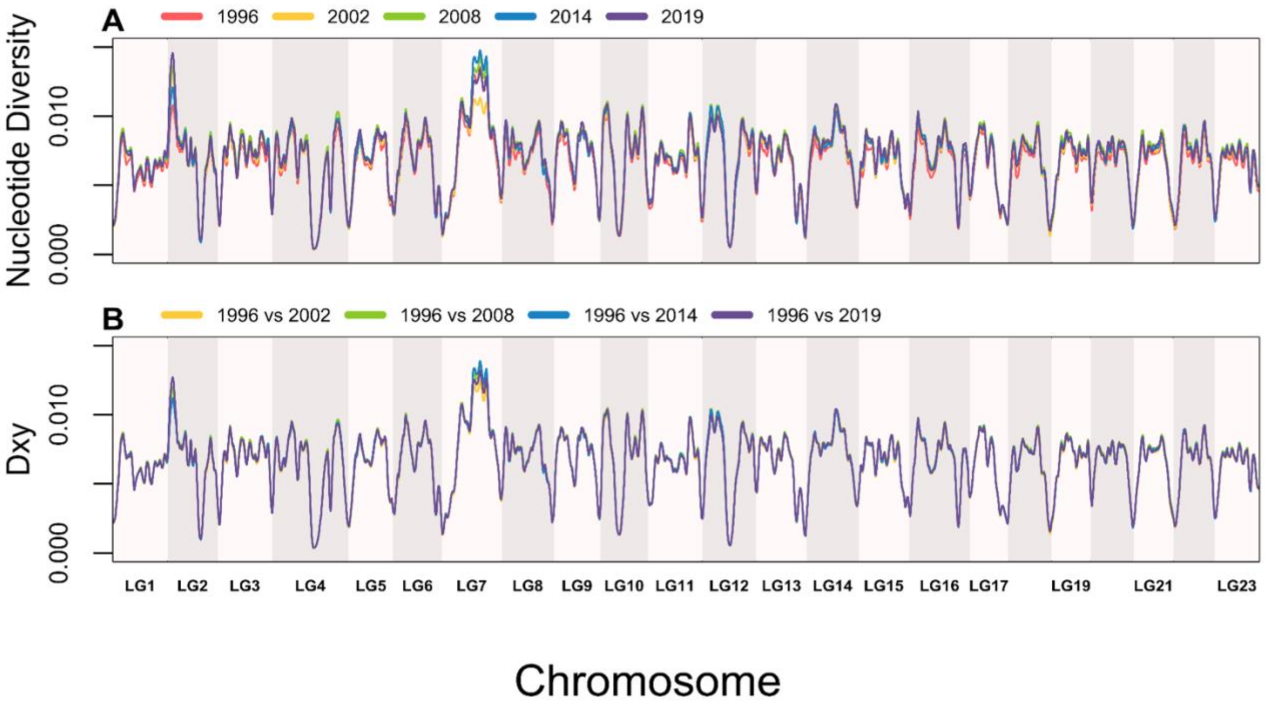

**Figure S7. Frequency distribution of lag-2 and lag-3 temporal covariance of randomly permuted 338 SNPs compared to GWA outliers.**

To detect signals of directional selection of GWA outliers which are correlated to growth performance, temporal covariance of allele frequency of the outliers were calculated and compared to values from randomly permuted 336 SNPs. The lag-2 and lag-3 time windows were used for calculating covariance to avoid shared time points, as in the inlets of each histogram visualised with a hypothetical allele frequency change.The frequency distribution of covariance of 1000 permutations of random SNPs are presented. The covariance was calculated in lag-2 time windows, namely cov(Δ1996-2008, Δ2002-2014) from top and cov(Δ2002-2014, Δ2008-2019), and lag-3 time window, cov(Δ1996-2014, Δ2002-2019) at the bottom. The red vertical solid lines depict the corresponding real covariance values of GWAS outlier SNPs.

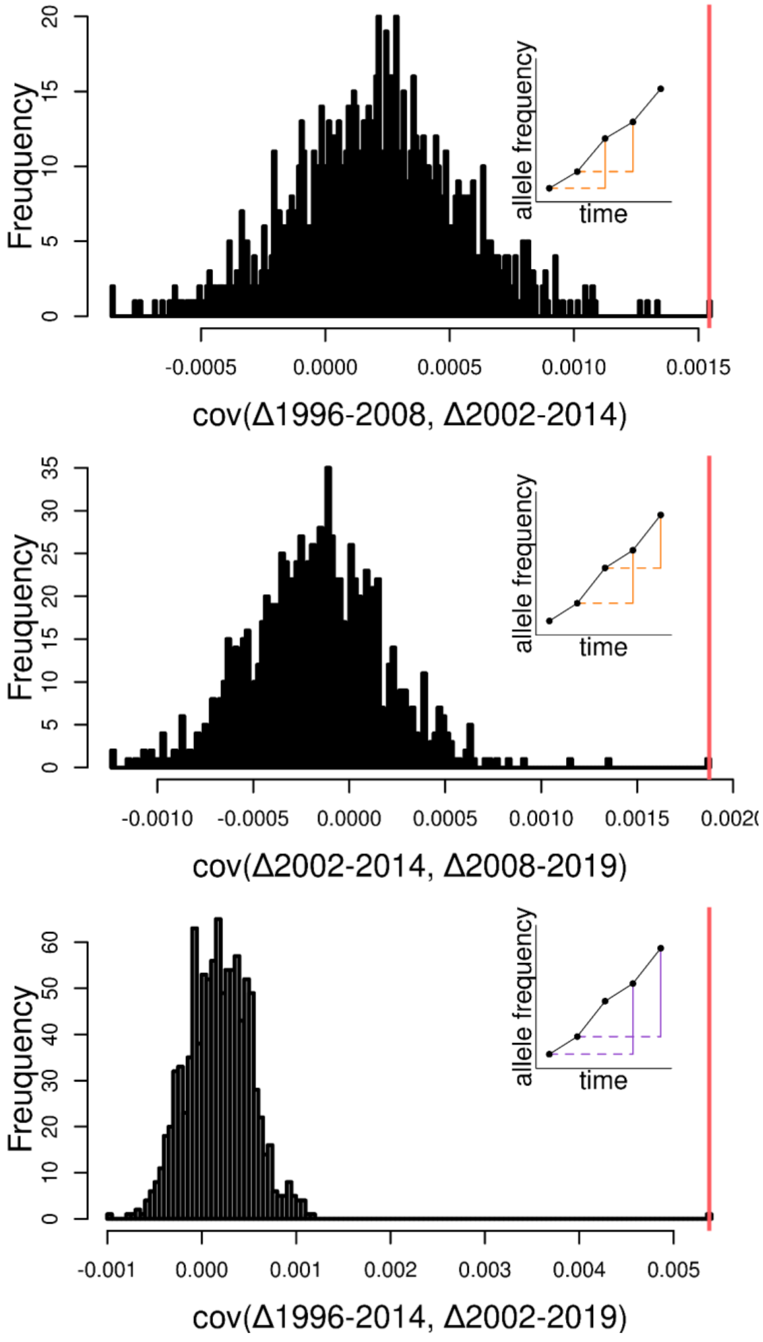

**Figure S8. Frequency distribution of the number of randomly overlapping windows of Fst and GWA outliers.**

A custom-made null model was used to assess the significance of the observed number of overlaps among Fst outlier windows and GWA outlier SNPs. To this end, 5000 random permutations of 336 SNPs were overlapped to $F_{st}$ outlier windows. When a GWA outlier SNP (or a randomly chosen SNP in a permutation) resides within a $F_{st}$ outlier window (20Kb), this window counts as an overlapping outlier window. The null distribution of the number of overlapping outlier windows is presented with the red line presenting the observed number of overlapping windows at 33 in real data.

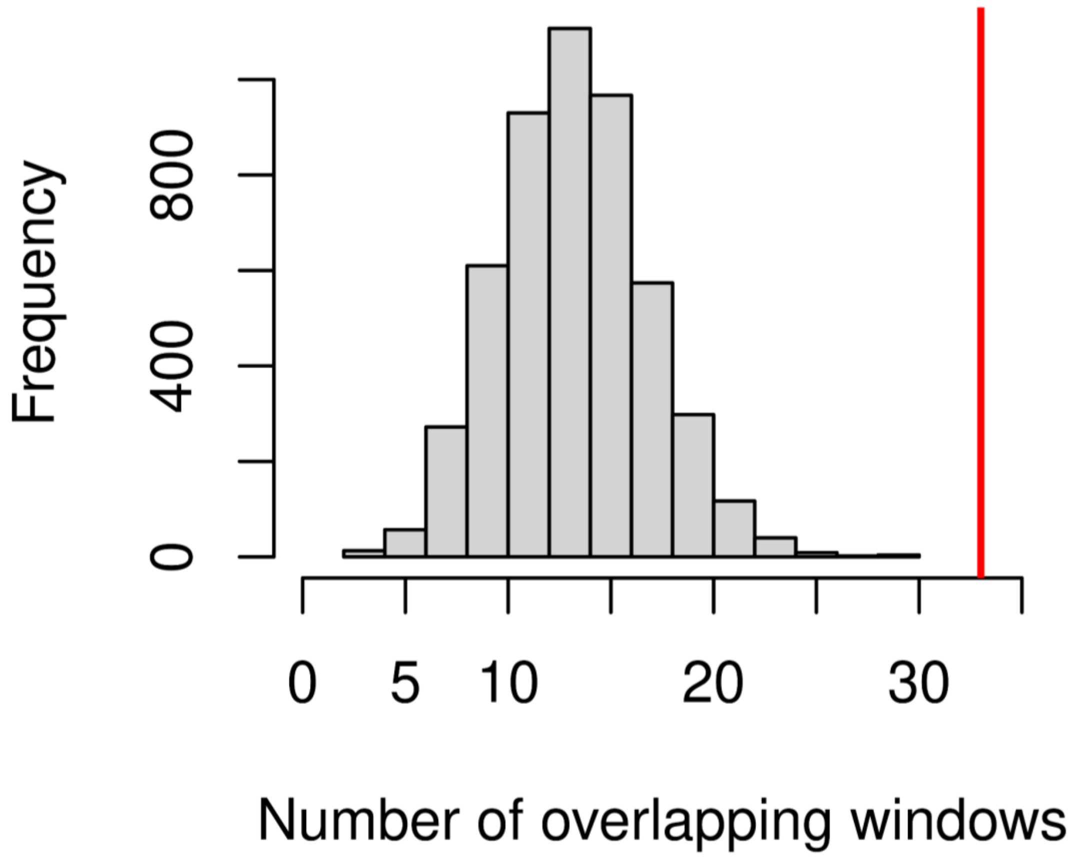

**Figure S9. Ancestry painting for inversion in LG12.**

To identify the double-crossover status of each individual, all “random” samples were examined by ancestry painting analysis, using four samples as reference of ancestral and derived homozygotes and two EBC as reference of double crossover. SNP sites between 6.5 and 7.5Mb within LG12 (reference to garMor3.0) which are fixed 80% in two sets of references were painted depending on the allelic status of ancestral or derived (green and yellow respectively as in legend), allowing for 20% of missingness in the data. The double crossover region is visually identifiable with higher frequency of derived alleles (marked with red vertical lines across all samples). With two rows of genomes in each individual, the genotype status was assigned in three sections, before, at and after the double crossover region. Sample names in the plot correspond to the sequencing names in Table S1.

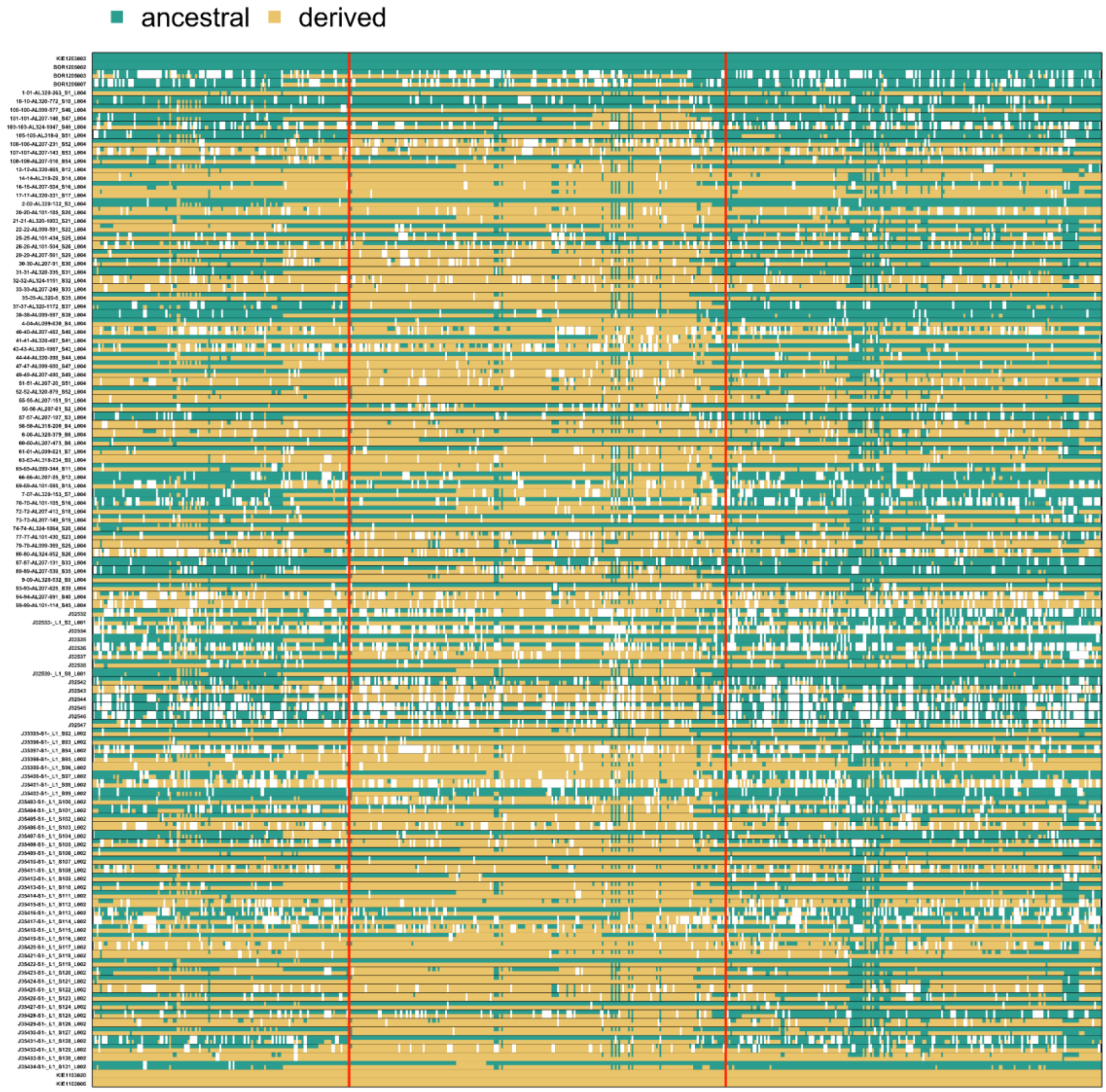

**Figure S10. Posterior distributions and model convergence of three Markov chain analyses to estimate von Bertalanffy parameters.**

The posterior distributions of four parameters used in the growth model (only three von growth parameters for all samples; *k.all, linf.all*, and *t0.all* and error *tol* are shown), in the left column, and corresponding trace plots of simulated three Markov chains over 100,000 iterations with a thinning rate of ten, in the right column, are plotted. Each chain starting from different initial values have converged adequately at the beginning of the iterations, indicating the robustness of model fit.

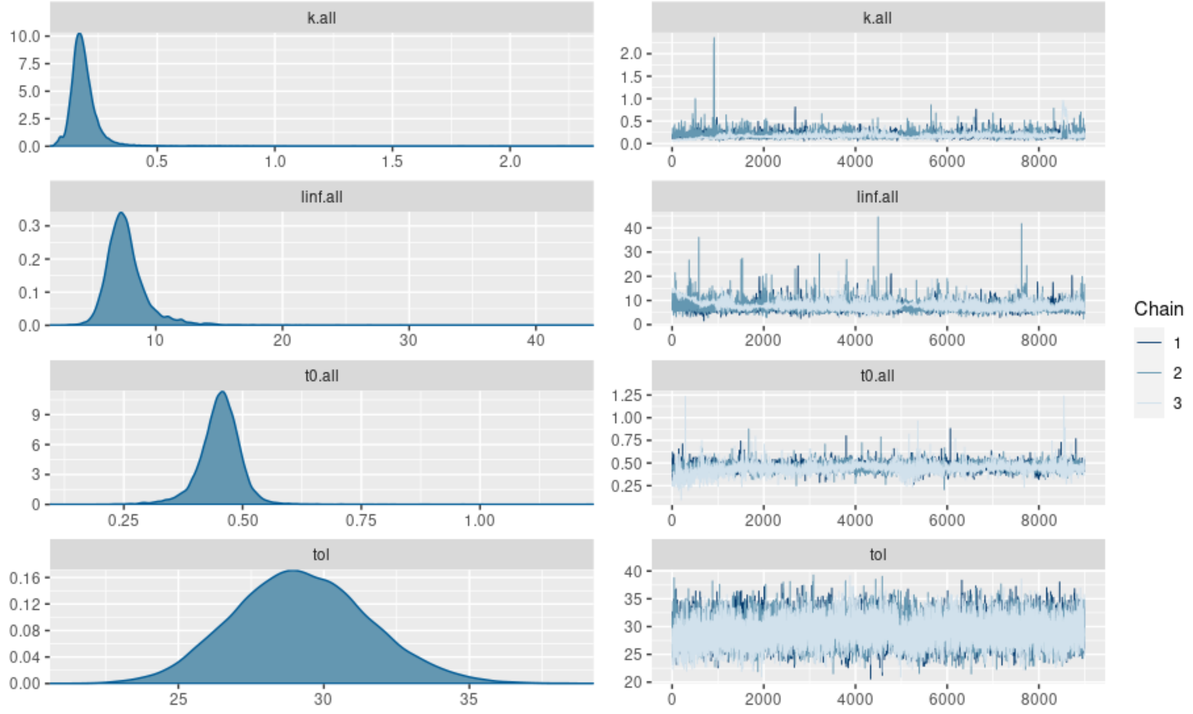

**Figure S11. Residuals of estimated otolith lengths for each age class.**

The residuals were calculated as (estimated otolith radii at age-observed otolith radii at age) / (observed otolith radii at age), matching the number of observations of each age class to assess the model fit. The discrepancy in variance of residuals in each age class is likely to come from the differences in the number of data points available in input (154 observations for age 1 and 1 observation for age 7).

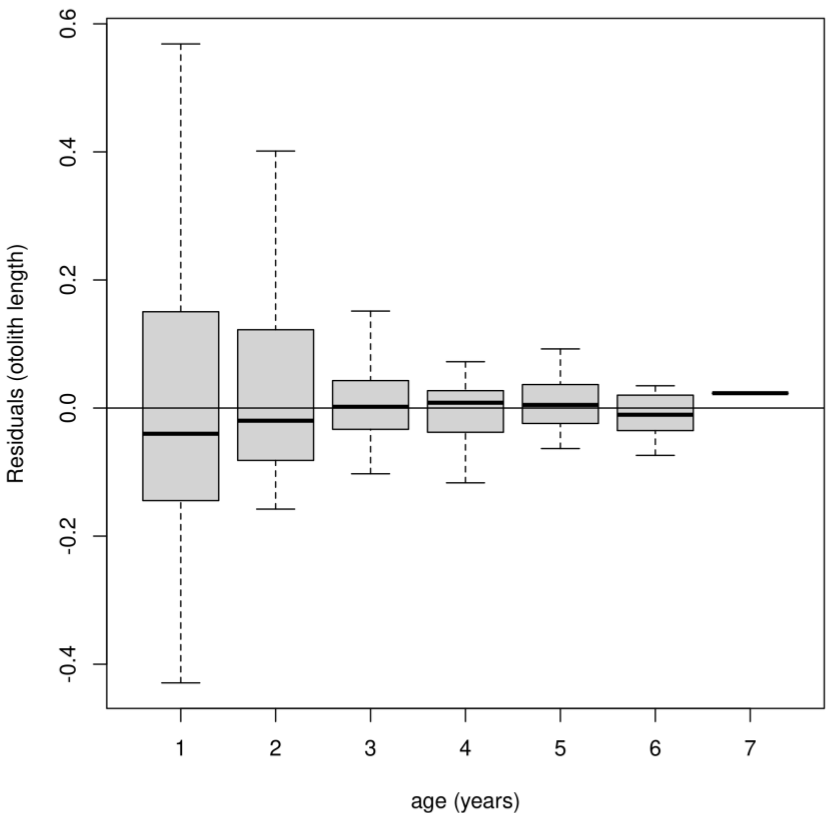

**Figure S12. Segregation and electrophoretic visualisation of microsatellite (MSAT) alleles in clean and contaminated samples.**

Four MSAT loci, (PGmo38 and Gmo5 blue; Tch11 black; GmoC18 red) were used to identify cross-contaminated samples. Prior knowledge of each MSAT length range in the resolution of 1bp was used to bin allele sizes using the GeneMarker® software. **A.** An example MSAT panel of a clean sample. Each locus shows two clear peaks, indicating a heterozygous status, marked with grey vertical lines. For example, in PGmo38 (left side of blue panel), a peak at 95bp and the other at 105bp appear as two different alleles in the locus. The smaller peaks before the highest peak in each allele is a typical noise pattern from a characteristic of MSAT analysis, called “stutter band”. The signal derives from incomplete products of multiplex PCR cycles, thus always smaller than the signal from complete products. **B.** An example of contaminated samples. When more than two individuals with different MSAT genotypes are mixed in the DNA samples, three or more peaks appear as shown. For example, in Gmo5 (right side of the blue panel), three peaks are identified. As stutter bands are always smaller and increasing towards the real peaks, the last peak at size 195 bp (marked in red box) is likely to be a real signal of an additional allele. The other MSAT sites also show more than three peaks even after considering stutter patterns. All MSAT sites of all individuals were examined manually to confirm and edit the automatic peak detection of the software. Samples with more than three peaks in at least one MSAT were excluded in the selection process.

**A**

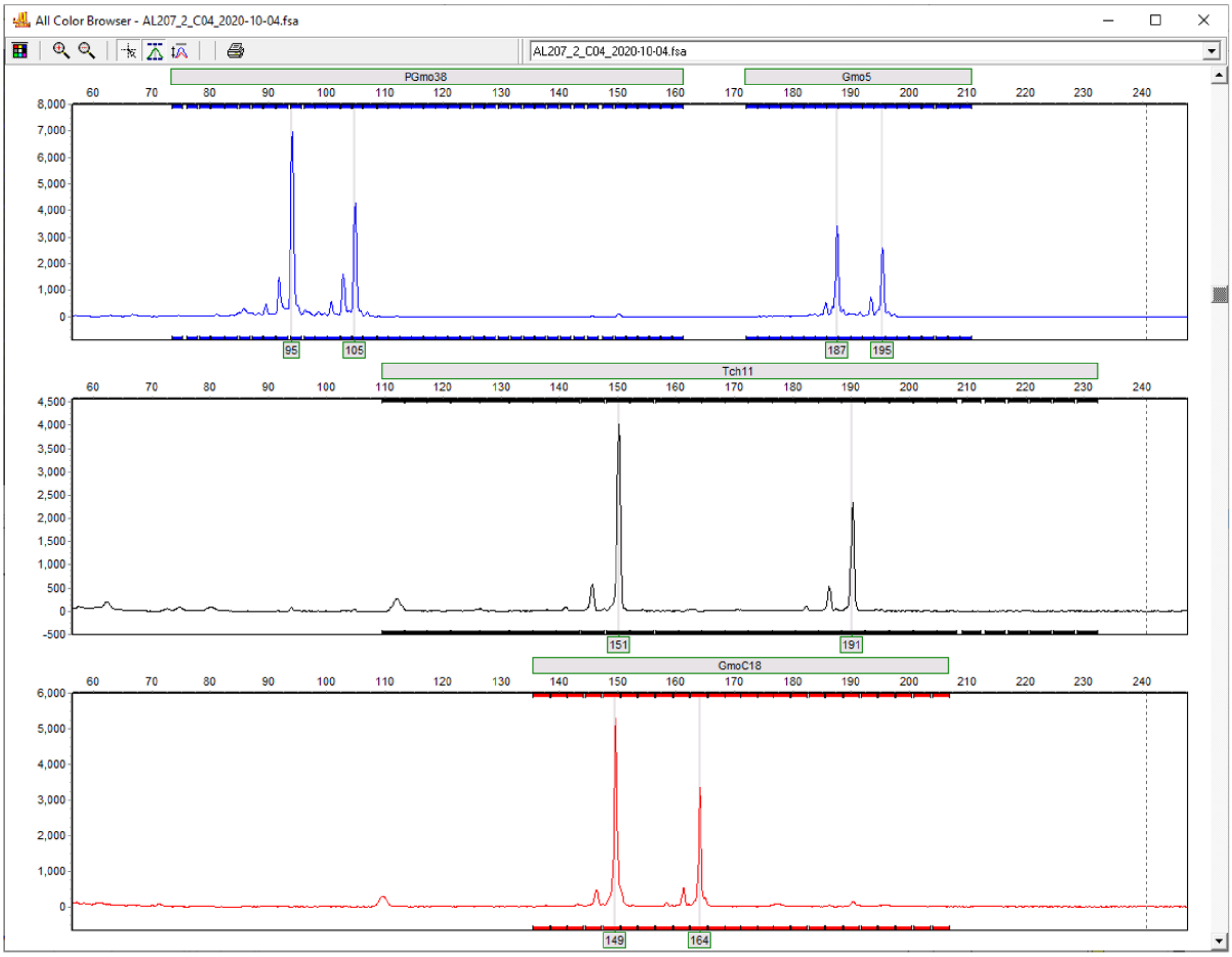

**B**

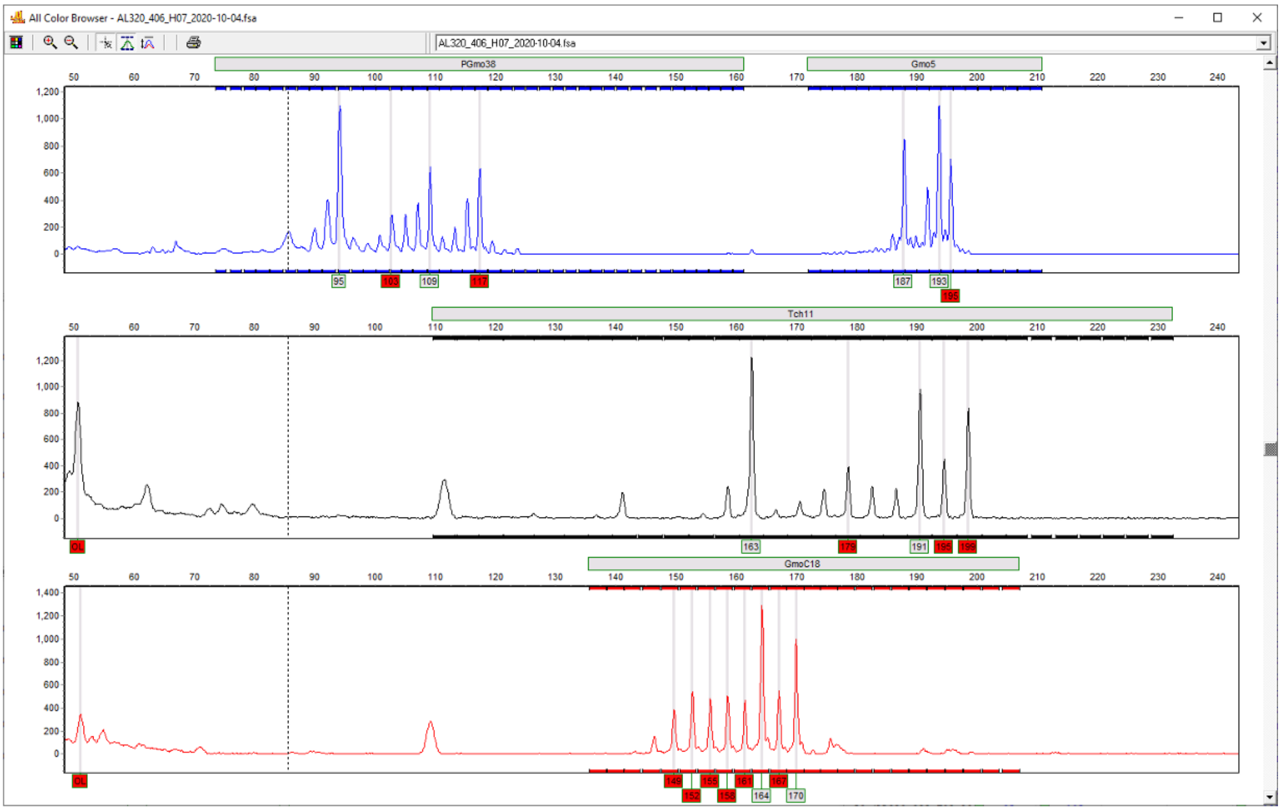
All Color Browser - AL320_406_H07_2020-10-04.fsa
AL320_406_H07_2020-10-04.fsa
PGmo38
Gmo5
Tch11
GmoC18
95
103
109
117
187
193
195
163
179
191
195
199
149
152
155
158
161
164
167
170

**Figure S13. Principal component analysis including cod individuals from Barth et al. 2019.**

Principal component analysis was done with all sequenced samples from this study, “random” and “phenotype” together with other Atlantic cod populations of the Baltic and North Sea from Barth et al. 2019, 23 EBC (BOR); 22 WBC (KIE); and 24 North Sea (NOR). This was to identify potential migrants in our samples and test for bias in sequencing. Nine individuals of “phenotype” samples which cluster with WBC individuals were excluded in any downstream analysis.

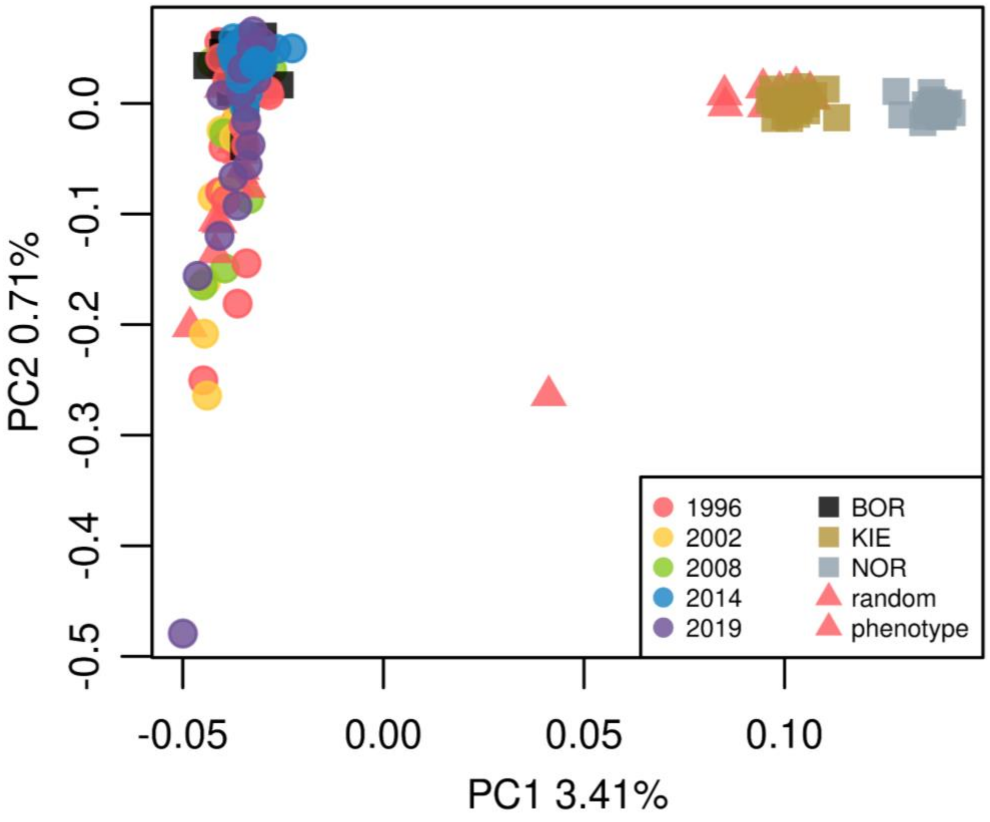

**Figure S14. Principal component analysis of 115 "random" samples.**

PCA was done with all variants of 4,685,343 SNPs (after filtering for MAF > 0.01). **A.** PC1 explains 2.25% of all variations in the genotypes and PC2 explains 1.57%. The clustering pattern observed disappears in Figure 1D once the SNPs within inverted regions in LG2, 7, and 12 are removed**. B.** Population structure on PC3 and PC4. Each individual is coded in colour according to the catch.

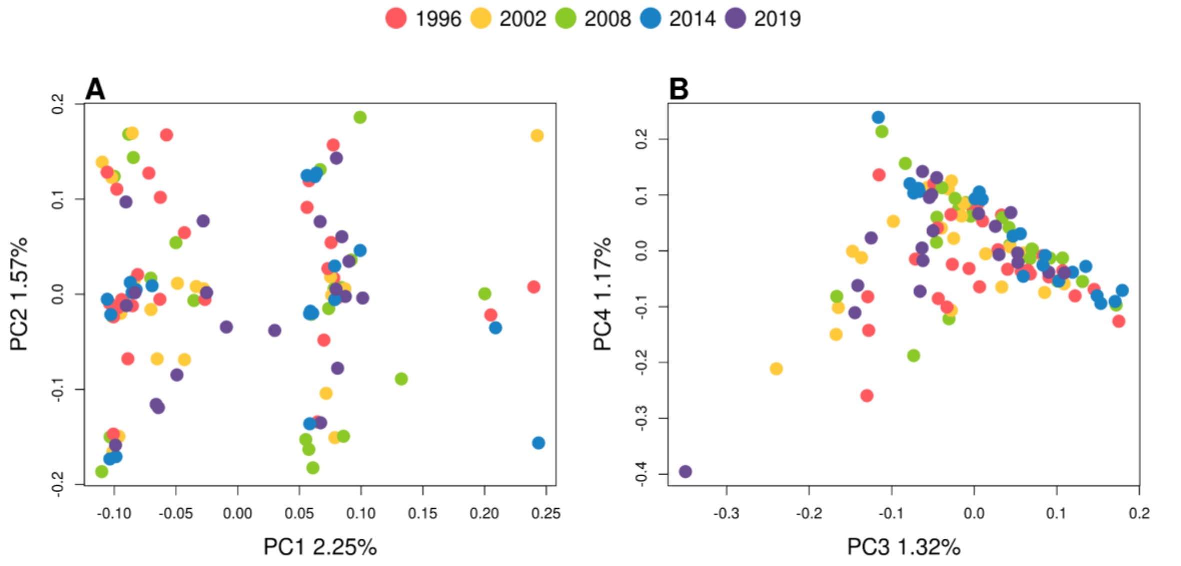

**Figure S15. QQ plot of expected and observed chi-squared p-values of the GWA.**

The observed chi-squared p-values of the GWA were plotted against chi-squared expected p-value distribution in a quantile-quantile (QQ) plot. The genomic inflation factor (= lambda), i.e., the median of chi-squared observed p-value divided by expected median of chi-squared p-value, was calculated. The overall fit of the expected versus the observed with a slight deviation at the highest end indicates an adequate correction of confounding factors during the GWA analysis.

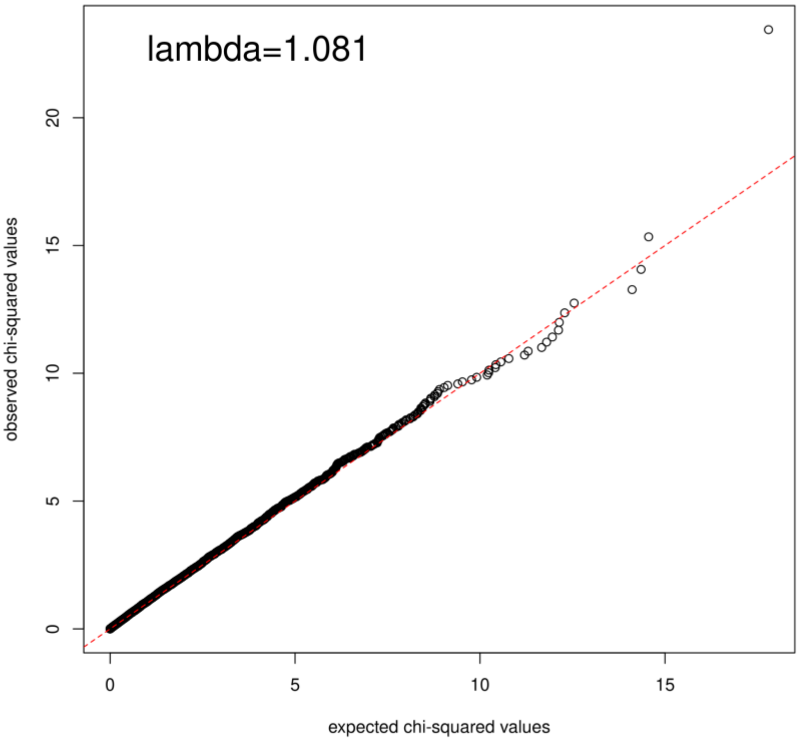

**Figure S16. Global and pairwise $F_{st}$ on groups of individuals based on the inversion status (LG2, LG7, and LG12).**

To determine the boundaries of inversion referenced to the reference genome gadMor3.0, $F_{st}$ values in 30Kb overlapping windows (in 15Kb steps) were computed using groups of individuals of different inversion status**.** After plotting the global $F_{st}$ values over the whole chromosome (row 1), approximate coordinates were chosen to zoom in (row 2) to decide the beginning and end of the inverted regions (row 3 and 5). The pairwise $F_{st}$ of two groups of homozygotes (ancestral and derived status) were also investigated to confirm the consistency of signals (row 4 and 6). Two locations were visually selected in the beginning and the end of inversions (Dashed vertical lines in row 3-6) to either include or exclude the regions depending on the downstream analysis conducted.

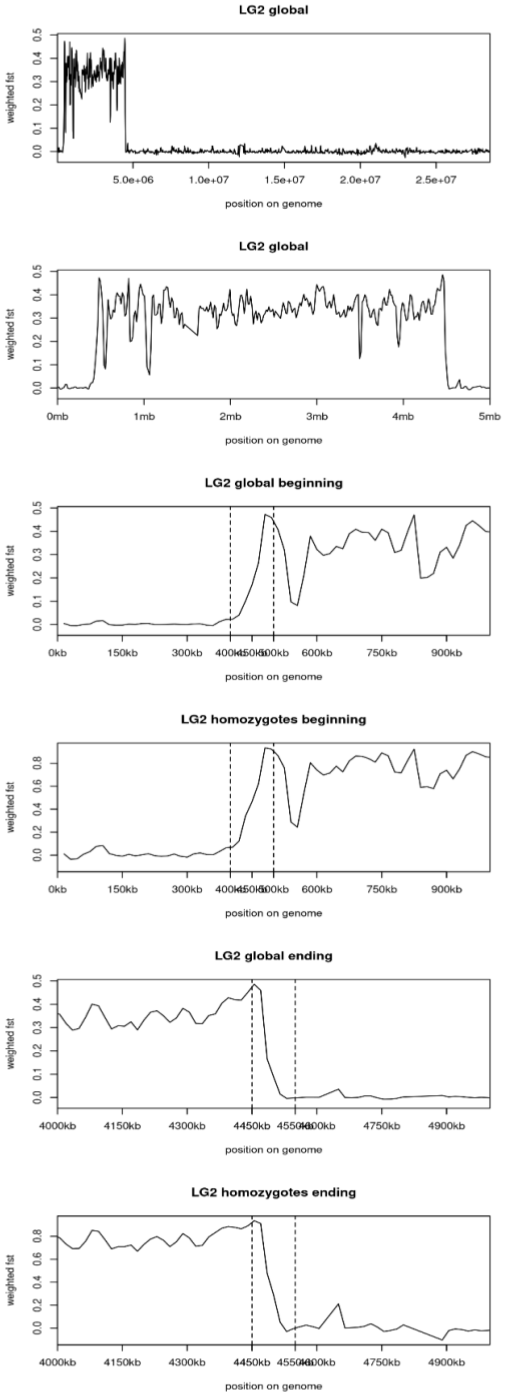

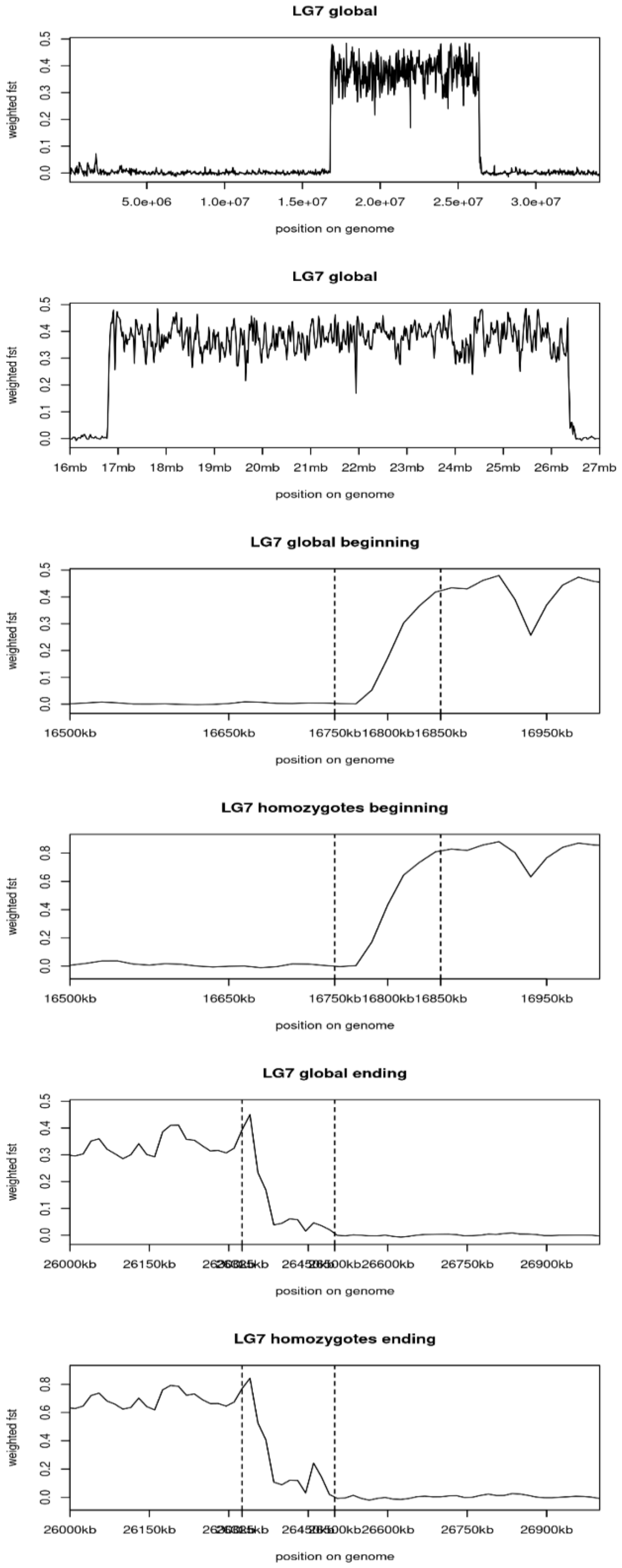
LG7 global
weighted fst
0.0 0.1 0.2 0.3 0.4 0.5
5.0e+06 1.0e+07 1.5e+07 2.0e+07 2.5e+07 3.0e+07
position on genome
LG7 global
weighted fst
0.0 0.1 0.2 0.3 0.4 0.5
16mb 17mb 18mb 19mb 20mb 21mb 22mb 23mb 24mb 25mb 26mb 27mb
position on genome
LG7 global beginning
weighted fst
0.0 0.1 0.2 0.3 0.4 0.5
16500kb 16650kb 16750kb 16800kb 16850kb 16950kb
position on genome
LG7 homozygotes beginning
weighted fst
0.0 0.2 0.4 0.6 0.8
16500kb 16650kb 16750kb 16800kb 16850kb 16950kb
position on genome
LG7 global ending
weighted fst
0.0 0.1 0.2 0.3 0.4 0.5
26000kb 26150kb 26600kb 26750kb 26900kb
position on genome
LG7 homozygotes ending
weighted fst
0.0 0.2 0.4 0.6 0.8
26000kb 26150kb 26600kb 26750kb 26900kb
position on genome

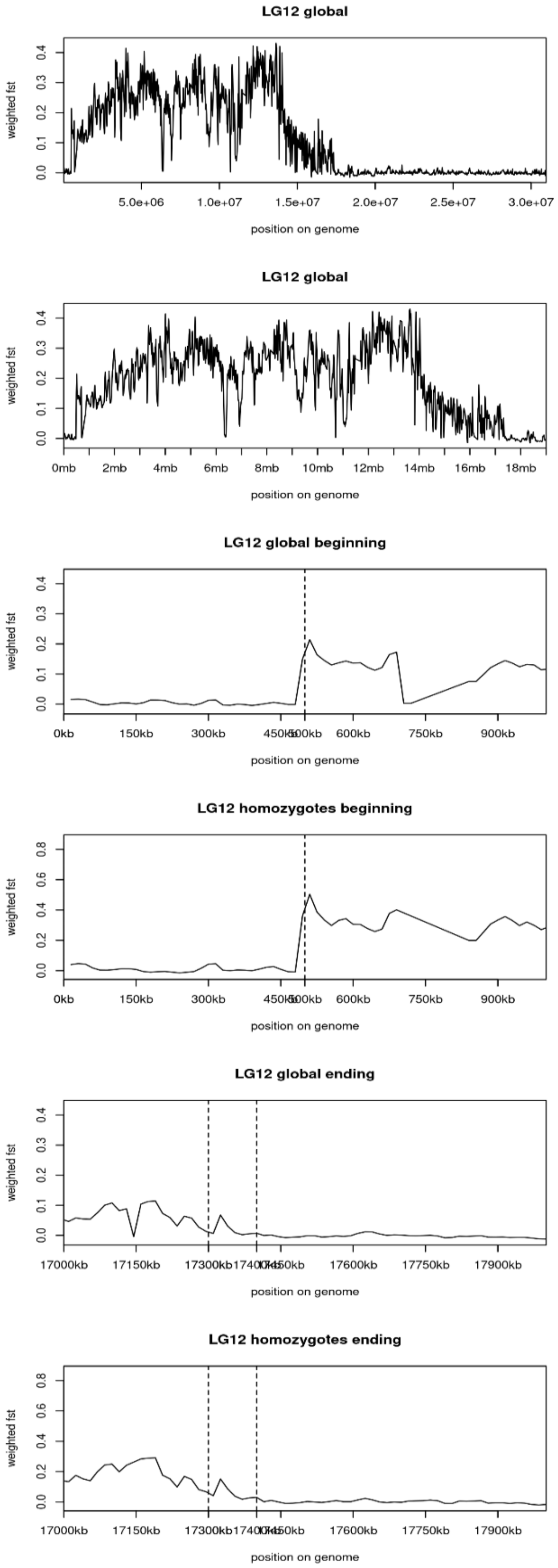
LG12 global
weighted fst
0.0 0.1 0.2 0.3 0.4
5.0e+06 1.0e+07 1.5e+07 2.0e+07 2.5e+07 3.0e+07
position on genome
LG12 global
weighted fst
0.0 0.1 0.2 0.3 0.4
0mb 2mb 4mb 6mb 8mb 10mb 12mb 14mb 16mb 18mb
position on genome
LG12 global beginning
weighted fst
0.0 0.1 0.2 0.3 0.4
0kb 150kb 300kb 450kb 500kb 600kb 750kb 900kb
position on genome
LG12 homozygotes beginning
weighted fst
0.0 0.2 0.4 0.6 0.8
0kb 150kb 300kb 450kb 500kb 600kb 750kb 900kb
position on genome
LG12 global ending
weighted fst
0.0 0.1 0.2 0.3 0.4
17000kb 17150kb 17300kb 17400kb 17450kb 17600kb 17750kb 17900kb
position on genome
LG12 homozygotes ending
weighted fst
0.0 0.2 0.4 0.6 0.8
17000kb 17150kb 17300kb 17400kb 17450kb 17600kb 17750kb 17900kb
position on genome

**Figure S17. Local principal component analysis on inverted chromosomal regions in LG02, LG07, and LG12 of the cod *Gadus morhua* genome.**

Local PCA was conducted using SNP sites within the inverted regions to identify individual inversion status. Three groups depending on the genotypes, ancestral and derived homozygotes on the sides and heterozygotes in the middle, are strongly clustered. Samples were assigned their status according to these clusters to calculate inversion frequency over time. Individuals are coloured according to the catch year.

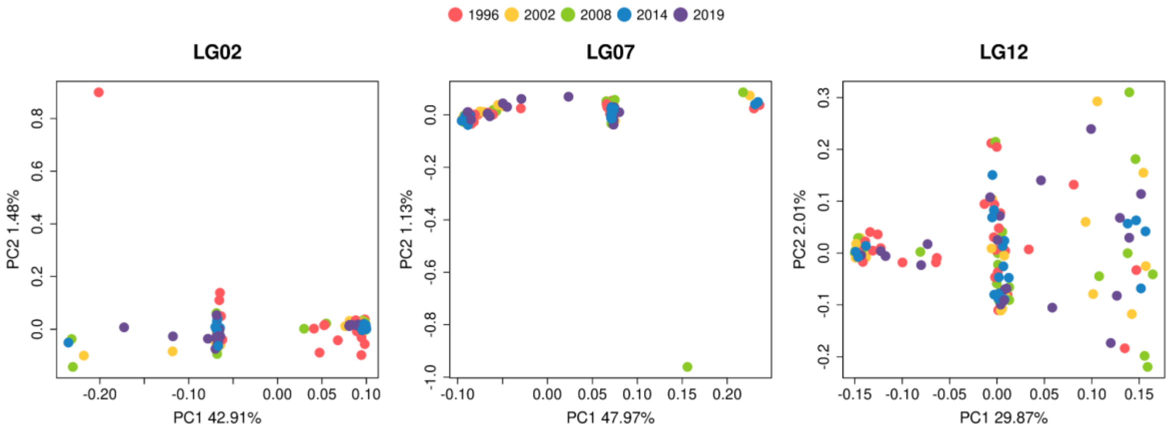

**Table S1. All samples used in this study and metadata.**

A total of 154 samples were subjected to sequencing and growth modelling. Two of them were excluded in any genetic analysis due to their low sequencing depth. "Otolith radius in mm" : the length from otolith core to the edge, "distance_min_x" : the distance from the core to the chemical annuli (minima) for each age estimate, "chemAge" : the estimated age from the age reading protocol, "sample.type" : the assignment of the sample type "random" and "phenotype" according to the sampling design. The samples were grouped into "bins" according to the catch year for growth modelling. The "phenotype" samples from 1996-1998 were grouped together with "random" samples from 1996 in "bin1". For all the genetic analysis, "SequencingName" were used and matched with "fishID" for metadata.

| fishID | catch.year | length in mm | sex | maturity | otolith radius in mm | chemAge | distance_min_1 | distance_min_2 | distance_min_3 | distance_min_4 | distance_min_5 | distance_min_6 | distance_min_7 | sample.type | bins | SequencingName |
|---|---|---|---|---|---|---|---|---|---|---|---|---|---|---|---|---|
| AL097_334 | 1996 | 420 | F | 1 | 3.211704 | 3 | 0.7882197 | 1.9999617 | 3.10577 | NA | NA | NA | NA | pheno | bin1 | 104-104-AL097-334_S50_L004 |
| AL097_430 | 1996 | 260 | M | 4 | 2.3528964 | 3 | 0.4396807879 | 1.319046909 | 2.270547 | NA | NA | NA | NA | pheno | bin1 | 75-75-AL097-430_S21_L004 |
| AL097_552 | 1996 | 410 | M | 1 | 3.30582 | 4 | 0.6611660184 | 1.385672 | 2.231428877 | 2.786335818 | NA | NA | NA | pheno | bin1 | 83-83-AL097-552_S29_L004 |
| AL099_189 | 1996 | 460 | F | 1 | 4.317564 | 7 | 0.8161973752 | 1.466789481 | 2.259326806 | 2.850775876 | 3.31210568 | NA | NA | pheno | bin1 | 39-39-AL099-189_S39_L004 |
| AL099_326 | 1996 | 270 | M | 5 | 2.1646641 | 3 | 0.4941093 | 1.423503 | 2.01172563 | NA | NA | NA | NA | pheno | bin1 | 64-64-AL099-326_S10_L004 |
| AL099_328 | 1996 | 400 | F | 1 | 2.9293566 | 3 | 0.5646945 | 2.1999582 | 2.70583023 | NA | NA | NA | NA | pheno | bin1 | 102-102-AL099-328_S48_L004 |
| AL099_344 | 1996 | 310 | M | 5 | 2.5175976 | 3 | 0.399993 | 1.2470361 | 2.35289643 | NA | NA | NA | NA | random | bin1 | 65-65-AL099-344_S11_L004 |
| AL099_35 | 1996 | 890 | F | 6 | 5.450875 | 5 | 0.49018495 | 1.39213 | 2.86269 | 4.27443 | 5.058725 | NA | NA | random | bin1 | J32532-_L1_S1_L002 |
| AL099_37 | 1996 | 1040 | F | 6 | 5.929299 | 6 | 0.70587 | 2.776419 | 3.6587559 | 4.5410901 | 5.1251896 | 5.7754189 | NA | random | bin1 | J32533-_L1_S2_L001 |
| AL099_577 | 1996 | 570 | F | 6 | 3.835218 | 3 | 0.7005473147 | 2.64784067 | 3.775258 | NA | NA | NA | NA | random | bin1 | 100-100-AL099-577_S46_L004 |
| AL099_591 | 1996 | 350 | M | 7 | 2.541129 | 4 | 0.4845875811 | 1.004636149 | 1.678331314 | 2.475258 | NA | NA | NA | random | bin1 | 22-22-AL099-591_S22_L004 |
| AL099_597 | 1996 | 530 | M | 6 | 4.258746 | 4 | 1.047039 | 2.294076 | 3.5411103 | 4.185647 | NA | NA | NA | random | bin1 | 38-38-AL099-597_S38_L004 |
| AL099_600 | 1996 | 340 | M | 6 | 3.199941 | 4 | 0.329406 | 1.2588003 | 2.105844 | 2.847006 | NA | NA | NA | random | bin1 | 47-47-AL099-600_S47_L004 |
| AL099_602 | 1996 | 270 | M | 6 | 2.4705438 | 3 | 0.7529295 | 1.2352734 | 1.5293817 | NA | NA | NA | NA | pheno | bin1 | 53-53-AL099-602_S53_L004 |

| | | | | | | | | | | | | | | | | |
|---|---|---|---|---|---|---|---|---|---|---|---|---|---|---|---|---|
| AL099_613 | 1996 | 580 | F | 6 | 3.92934 | 3 | 0.623517 | 2.4705417 | 3.450258 | NA | NA | NA | NA | random | bin1 | J32535-_L1_S3_L002 |
| AL099_621 | 1996 | 630 | M | 7 | 4.599915 | 6 | 0.4138730706 | 1.064247408 | 2.38273266 | 3.068580604 | 3.783991621 | 4.245165247 | NA | random | bin1 | 61-61-AL099-621_S7_L004 |
| AL099_630 | 1996 | 430 | M | 6 | 3.517578 | 3 | 0.5665854198 | 1.085960329 | 2.679497306 | NA | NA | NA | NA | random | bin1 | 4-04-AL099-630_S4_L004 |
| AL099_636 | 1996 | 620 | M | 7 | 4.8430445 | 5 | 0.6666505 | 1.1764455 | 2.1372095 | 3.1450465 | 4.0783495 | NA | NA | random | bin1 | J32536-_L1_S4_L002 |
| AL099_78 | 1996 | 560 | M | 6 | 4.411681 | 6 | 0.45097045 | 1.1176255 | 1.68624 | 2.4901455 | 3.313661 | 3.7058095 | NA | random | bin1 | J32534-_L1_S2_L002 |
| AL101_105 | 1996 | 570 | F | 7 | 4.623444 | 5 | 0.6015187318 | 2.193776675 | 3.290665614 | 3.915776082 | 4.550167 | NA | NA | random | bin1 | 70-70-AL101-105_S16_L004 |
| AL101_114 | 1996 | 580 | F | 7 | 4.599912 | 6 | 0.823515 | 1.435269 | 2.141136 | 2.8411237 | 3.646989 | 4.5 | NA | random | bin1 | 99-99-AL101-114_S45_L004 |
| AL101_185 | 1996 | 590 | M | 6 | 4.282269 | 6 | 0.4507659285 | 1.328570843 | 2.052166361 | 2.882524952 | 3.6298459 | 4.2 | NA | random | bin1 | 20-20-AL101-185_S20_L004 |
| AL101_214 | 1996 | 540 | F | 8 | 3.164646 | 4 | 0.6978097159 | 1.564543397 | 2.323111806 | 3.1 | NA | NA | NA | random | bin1 | J32537-_L1_S5_L002 |
| AL101_287 | 1996 | 460 | M | 7 | 3.682284 | 4 | 0.482346 | 0.988218 | 2.5058343 | 3.6 | NA | NA | NA | random | bin1 | J32538-_L1_S6_L002 |
| AL101_288 | 1996 | 550 | F | 6 | 4.388151 | 7 | 0.5946012489 | 1.117850348 | 1.95028967 | 2.473538466 | 3.24652009 | 3.853009118 | 4.102741642 | random | bin1 | J32539-_L1_S8_L001 |
| AL101_344 | 1996 | 430 | F | 7 | 3.670518 | 4 | 1.1411547 | 1.741143 | 2.8352406 | 3.6 | NA | NA | NA | random | bin1 | J32542-_L1_S9_L002 |
| AL101_422 | 1996 | 560 | F | 7 | 3.905811 | 5 | 0.835278 | 1.470564 | 2.1764283 | 2.7528929 | 3.71757858 | NA | NA | random | bin1 | J32543-_L1_S10_L002 |
| AL101_423 | 1996 | 480 | F | 7 | 3.435228 | 3 | 0.8470425 | 1.8940806 | 3.24699558 | NA | NA | NA | NA | random | bin1 | J32544-_L1_S11_L002 |
| AL101_430 | 1996 | 540 | F | 7 | 4.25874 | 6 | 0.5323416666 | 1.005538617 | 1.940091808 | 2.489621822 | 3.111246793 | 3.927502105 | NA | random | bin1 | 77-77-AL101-430_S23_L004 |
| AL101_434 | 1996 | 430 | M | 7 | 2.9175936 | 5 | 0.462548792 | 1.079270837 | 1.897623088 | 2.039940722 | 2.407611293 | NA | NA | random | bin1 | 25-25-AL101-434_S25_L004 |
| AL101_442 | 1996 | 800 | F | 7 | 5.305779 | 6 | 0.6735619839 | 1.890701522 | 2.812416969 | 3.521431169 | 3.94683969 | 4.774019609 | NA | random | bin1 | J32545-_L1_S12_L002 |
| AL101_504 | 1996 | 460 | F | 7 | 3.70581 | 4 | 0.5701255101 | 1.045227073 | 2.316130644 | 3.539523029 | NA | NA | NA | random | bin1 | 26-26-AL101-504_S26_L004 |
| AL101_523 | 1996 | 650 | F | 7 | 4.599912 | 5 | 0.6148993755 | 1.288922183 | 2.636967799 | 3.748516972 | 4.4 | NA | NA | random | bin1 | J32546-_L1_S13_L002 |
| AL101_546 | 1996 | 500 | M | 1 | 3.449085 | 4 | 0.6470439 | 1.5058527 | 2.6843898 | 3.28437795 | NA | NA | NA | pheno | bin1 | 76-76-AL101-546_S22_L004 |
| AL101_595 | 1996 | 470 | M | 7 | 3.352878 | 4 | 0.2833422798 | 1.097950431 | 2.184092526 | 3.093147742 | NA | NA | NA | random | bin1 | 69-69-AL101-595_S15_L004 |
| AL101_623 | 1996 | 220 | M | 6 | 2.235252 | 3 | 0.3882306 | 1.1764512 | 1.8940839 | NA | NA | NA | NA | pheno | bin1 | 95-95-AL101-623_S41_L004 |

| | | | | | | | | | | | | | | | | |
|---|---|---|---|---|---|---|---|---|---|---|---|---|---|---|---|---|
| AL101_679 | 1996 | 510 | F | 7 | 3.4313135 | 4 | 0.2941162 | 1.2744875 | 2.0391795 | 2.8234765 | NA | NA | NA | random | bin1 | J32547-_L1_S14_L002 |
| AL111_235 | 1997 | 270 | M | 5 | 2.3293683 | 2 | 0.7645091697 | 2.126296225 | NA | NA | NA | NA | NA | pheno | bin1 | 81-81-AL111-235_S27_L004 |
| AL111_28 | 1997 | 430 | F | 1 | 3.423462 | 4 | 0.5212224352 | 1.149051534 | 2.025651062 | 3.008856348 | NA | NA | NA | pheno | bin1 | 67-67-AL111-28_S13_L004 |
| AL114_826 | 1997 | 340 | F | 5 | 3.30582 | 3 | 0.8649636474 | 2.322368675 | 3.2 | NA | NA | NA | NA | pheno | bin1 | 13-13-AL114-826_S13_L004 |
| AL114_868 | 1997 | 350 | F | 4 | 3.246996 | 3 | 0.8176741991 | 2.287117683 | 3.128492023 | NA | NA | NA | NA | pheno | bin1 | 5-05-AL114-868_S5_L004 |
| AL114_937 | 1997 | 350 | F | 6 | 3.376407 | 3 | 0.8487897867 | 2.416796168 | 3.2 | NA | NA | NA | NA | pheno | bin1 | 45-45-AL114-937_S45_L004 |
| AL129_106 | 1998 | 400 | F | 1 | 2.9528847 | 4 | 0.399993 | 1.4823252 | 2.2234863 | 2.76465522 | NA | NA | NA | pheno | bin1 | 18-18-AL129-106_S18_L004 |
| AL129_148 | 1998 | 350 | F | 4 | 2.778504 | 5 | 0.4823439 | 1.0470411 | 1.623498 | 2.0726367 | 2.48439267 | NA | NA | pheno | bin1 | 19-19-AL129-148_S19_L004 |
| AL129_243 | 1998 | 240 | M | 6 | 2.1528984 | 2 | 0.4163059375 | 1.760383449 | NA | NA | NA | NA | NA | pheno | bin1 | 42-42-AL129-243_S42_L004 |
| AL129_266 | 1998 | 300 | F | 4 | 2.5293633 | 3 | 0.5343765927 | 1.033124708 | 1.971240748 | NA | NA | NA | NA | pheno | bin1 | 50-50-AL129-266_S50_L004 |
| AL129_339 | 1998 | 400 | F | 1 | 2.9881758 | 3 | 0.8267653782 | 1.759833805 | 2.326756389 | NA | NA | NA | NA | pheno | bin1 | 15-15-AL129-339_S15_L004 |
| AL129_369 | 1998 | 350 | F | 4 | 2.8705353 | 3 | 0.5551164224 | 1.665348665 | 2.23227125 | NA | NA | NA | NA | random | bin1 | 23-23-AL129-369_S23_L004 |
| AL129_377 | 1998 | 410 | M | 1 | 3.15288 | 5 | 0.611754 | 1.1646843 | 1.729377 | 2.388189 | 2.91759021 | NA | NA | pheno | bin1 | 62-62-AL129-377_S8_L004 |
| AL129_389 | 1998 | 410 | F | 1 | 2.8352403 | 2 | 0.4016601013 | 2.043735052 | NA | NA | NA | NA | NA | pheno | bin1 | 3-03-AL129-389_S3_L004 |
| AL129_401 | 1998 | 410 | F | 1 | 2.9764101 | 2 | 0.6640568246 | 2.430933466 | NA | NA | NA | NA | NA | pheno | bin1 | 90-90-AL129-401_S36_L004 |
| AL129_419 | 1998 | 270 | M | 6 | 2.4234843 | 3 | 0.5438058898 | 1.525022739 | 2.116114997 | NA | NA | NA | NA | pheno | bin1 | 86-86-AL129-419_S32_L004 |
| AL129_467 | 1998 | 270 | M | 6 | 2.2470147 | 3 | 0.5468895363 | 1.47423065 | 2.033013177 | NA | NA | NA | NA | pheno | bin1 | 36-36-AL129-467_S36_L004 |
| AL129_47 | 1998 | 400 | F | 1 | 3.30582 | 4 | 0.6493552721 | 1.157034585 | 2.160588787 | 2.739108187 | NA | NA | NA | pheno | bin1 | 28-28-AL129-47_S28_L004 |
| AL129_87 | 1998 | 350 | F | 4 | 2.8117119 | 2 | 0.8779157876 | 2.325297341 | NA | NA | NA | NA | NA | pheno | bin1 | 34-34-AL129-87_S34_L004 |
| AL130_199 | 1998 | 260 | M | 6 | 2.5528887 | 3 | 0.5964681468 | 1.049787613 | 2.290440306 | NA | NA | NA | NA | pheno | bin1 | 98-98-AL130-199_S44_L004 |
| AL130_206 | 1998 | 230 | M | 6 | 2.5293633 | 2 | 0.9928371908 | 2.115683232 | NA | NA | NA | NA | NA | pheno | bin1 | 88-88-AL130-206_S34_L004 |
| AL130_223 | 1998 | 420 | F | 1 | 3.176409 | 3 | 0.7851854841 | 2.010541617 | 2.938476416 | NA | NA | NA | NA | pheno | bin1 | 8-08-AL130-223_S8_L004 |

| | | | | | | | | | | | | | | | | |
|---|---|---|---|---|---|---|---|---|---|---|---|---|---|---|---|---|
| AL130_244 | 1998 | 330 | F | 6 | 2.9881803 | 2 | 0.9249138092 | 2.027694449 | NA | NA | NA | NA | NA | pheno | bin1 | 78-78-AL130-244_S24_L004 |
| AL130_429 | 1998 | 410 | F | 1 | 3.129351 | 3 | 0.4978527531 | 1.967698629 | 2.939692628 | NA | NA | NA | NA | pheno | bin1 | 48-48-AL130-429_S48_L004 |
| AL130_440 | 1998 | 520 | F | 1 | 3.576402 | 3 | 0.6750192331 | 1.78820314 | 3.398767277 | NA | NA | NA | NA | pheno | bin1 | 91-91-AL130-440_S37_L004 |
| AL130_68 | 1998 | 420 | F | 1 | 3.23523 | 3 | 0.8549463871 | 2.113527618 | 2.680282951 | NA | NA | NA | NA | pheno | bin1 | 46-46-AL130-68_S46_L004 |
| AL133_100 | 1998 | 450 | F | 1 | 3.552873 | 3 | 0.8882189846 | 2.214623884 | 3.173899831 | NA | NA | NA | NA | pheno | bin1 | 97-97-AL133-100_S43_L004 |
| AL133_116 | 1998 | 340 | F | 6 | 2.9528862 | 3 | 0.6522451519 | 1.482374658 | 2.644552459 | NA | NA | NA | NA | pheno | bin1 | 27-27-AL133-116_S27_L004 |
| AL133_119 | 1998 | 320 | F | 6 | 2.6117169 | 3 | 0.6522451519 | 1.197758119 | 2.644552459 | NA | NA | NA | NA | pheno | bin1 | 71-71-AL133-119_S17_L004 |
| AL133_136 | 1998 | 470 | F | 1 | 3.399936 | 4 | 0.8854023731 | 1.605527096 | 2.679811202 | 3.175636356 | NA | NA | NA | pheno | bin1 | 24-24-AL133-136_S24_L004 |
| AL207_131 | 2002 | 390 | M | 7 | 2.9881758 | 5 | 0.6284651135 | 1.209496947 | 1.968402221 | 2.407140865 | 2.845879599 | NA | NA | random | bin2 | 87-87-AL207-131_S33_L004 |
| AL207_143 | 2002 | 410 | F | 7 | 2.9293536 | 4 | 0.3543567045 | 1.098505905 | 1.759972496 | 2.64036229 | NA | NA | NA | random | bin2 | 107-107-AL207-143_S53_L004 |
| AL207_146 | 2002 | 440 | F | 7 | 3.764634 | 4 | 0.694104 | 1.3882092 | 2.5528917 | 3.2 | NA | NA | NA | random | bin2 | 101-101-AL207-146_S47_L004 |
| AL207_149 | 2002 | 520 | F | 6 | 4.035219 | 6 | 0.5427495625 | 0.8731188615 | 1.392270316 | 2.312582686 | 3.138505331 | 3.763845804 | NA | random | bin2 | 73-73-AL207-149_S19_L004 |
| AL207_151 | 2002 | 640 | F | 6 | 4.117569 | 4 | 0.741164 | 2.164665 | 3.1411167 | 3.87051285 | NA | NA | NA | random | bin2 | 55-55-AL207-151_S1_L004 |
| AL207_197 | 2002 | 640 | M | 9 | 4.211685 | 4 | 0.599988 | 1.5 | 2.5764198 | 3.4822851 | NA | NA | NA | random | bin2 | 57-57-AL207-197_S3_L004 |
| AL207_20 | 2002 | 440 | F | 7 | 3.364644 | 4 | 0.5999913 | 1.6587927 | 2.270547 | 3 | NA | NA | NA | random | bin2 | 51-51-AL207-20_S51_L004 |
| AL207_231 | 2002 | 390 | F | 7 | 3.517578 | 4 | 0.7224642069 | 1.871302639 | 2.534550964 | 3.387294944 | NA | NA | NA | random | bin2 | 106-106-AL207-231_S52_L004 |
| AL207_249 | 2002 | 420 | M | 7 | 3.69405 | 4 | 0.647049 | 1.3646817 | 2.5999524 | 3.44699532 | NA | NA | NA | random | bin2 | 33-33-AL207-249_S33_L004 |
| AL207_25 | 2002 | 340 | F | 8 | 2.91759 | 3 | 0.858807 | 1.8117303 | 2.68230021 | NA | NA | NA | NA | random | bin2 | 66-66-AL207-25_S12_L004 |
| AL207_413 | 2002 | 430 | F | 7 | 3.894045 | 5 | 0.7338329108 | 1.810910549 | 2.603922628 | 3.065524183 | 3.633654307 | NA | NA | random | bin2 | 72-72-AL207-413_S18_L004 |
| AL207_462 | 2002 | 340 | M | 7 | 3.317583 | 4 | 0.6611543444 | 1.711921929 | 2.59739611 | 3.187715182 | NA | NA | NA | random | bin2 | 40-40-AL207-462_S40_L004 |
| AL207_473 | 2002 | 280 | M | 7 | 2.3764248 | 3 | 0.532034494 | 1.265062102 | 2.199079989 | NA | NA | NA | NA | random | bin2 | 60-60-AL207-473_S6_L004 |
| AL207_490 | 2002 | 340 | M | 6 | 2.9432226 | 3 | 0.8782172608 | 1.685232469 | 2.136207452 | NA | NA | NA | NA | random | bin2 | 49-49-AL207-490_S49_L004 |

| | | | | | | | | | | | | | | | | |
|---|---|---|---|---|---|---|---|---|---|---|---|---|---|---|---|---|
| AL207_504 | 2002 | 300 | M | 7 | 2.7646581 | 4 | 0.5220840928 | 1.340798843 | 1.791689375 | 2.444288742 | NA | NA | NA | random | bin2 | 16-16-AL207-504_S16_L004 |
| AL207_516 | 2002 | 410 | F | 5 | 3.211704 | 4 | 0.7058661 | 1.788198 | 2.2823076 | 3.01170594 | NA | NA | NA | random | bin2 | 108-108-AL207-516_S54_L004 |
| AL207_539 | 2002 | 370 | M | 7 | 2.9999472 | 4 | 0.3188930824 | 1.027539658 | 1.99602551 | 2.740107833 | NA | NA | NA | random | bin2 | 89-89-AL207-539_S35_L004 |
| AL207_561 | 2002 | 320 | M | 7 | 3.69405 | 4 | 0.6941085 | 1.2588 | 2.4352512 | 3.50581758 | NA | NA | NA | random | bin2 | 29-29-AL207-561_S29_L004 |
| AL207_61 | 2002 | 360 | F | 7 | 3.376407 | 4 | 0.5194484002 | 1.35 | 2.184042542 | 3.104879284 | NA | NA | NA | random | bin2 | 56-56-AL207-61_S2_L004 |
| AL207_625 | 2002 | 410 | M | 7 | 3.752868 | 4 | 0.3433214951 | 2.048094775 | 3.042544295 | 3.634480872 | NA | NA | NA | random | bin2 | 93-93-AL207-625_S39_L004 |
| AL207_691 | 2002 | 270 | M | 6 | 2.36466 | 2 | 0.9340402522 | 1.927195885 | NA | NA | NA | NA | NA | random | bin2 | 94-94-AL207-691_S40_L004 |
| AL207_91 | 2002 | 300 | M | 8 | 2.5528917 | 4 | 0.5936973173 | 1.270509752 | 1.733591673 | 2.256044477 | NA | NA | NA | random | bin2 | 30-30-AL207-91_S30_L004 |
| AL318_200 | 2008 | 350 | M | 5 | 3.188178 | 4 | 0.3081526075 | 0.687416774 | 1.481498318 | 2.69039754 | NA | NA | NA | random | bin3 | 58-58-AL318-200_S4_L004 |
| AL318_234 | 2008 | 350 | F | 5 | 3.258759 | 6 | 0.5925007683 | 1.007250974 | 1.824903442 | 2.322606469 | 2.784755696 | 3.2 | NA | random | bin3 | 63-63-AL318-234_S9_L004 |
| AL318_29 | 2008 | 420 | F | 5 | 3.905808 | 6 | 0.270582 | 0.599988 | 1.8470214 | 2.541126 | 3.352875 | 3.9 | NA | random | bin3 | 14-14-AL318-29_S14_L004 |
| AL318_9 | 2008 | 240 | M | 5 | 2.2587816 | 3 | 0.4139116985 | 0.9697372029 | 1.785736611 | NA | NA | NA | NA | random | bin3 | 105-105-AL318-9_S51_L004 |
| AL320_1003 | 2008 | 480 | F | 5 | 3.517581 | 4 | 0.3316183982 | 1.136999585 | 2.250310503 | 3.3 | NA | NA | NA | random | bin3 | 21-21-AL320-1003_S21_L004 |
| AL320_1067 | 2008 | 310 | M | 5 | 2.847 | 3 | 0.8778179102 | 1.51839758 | 2.669068386 | NA | NA | NA | NA | random | bin3 | 43-43-AL320-1067_S43_L004 |
| AL320_1172 | 2008 | 310 | M | 7 | 2.8352403 | 3 | 0.7473292852 | 1.969170141 | 2.6 | NA | NA | NA | NA | random | bin3 | 37-37-AL320-1172_S37_L004 |
| AL320_132 | 2008 | 490 | M | 5 | 4.117566 | 5 | 0.3775425303 | 1.156218583 | 2.465818032 | 3.374278542 | 3.9 | NA | NA | random | bin3 | 2-02-AL320-132_S2_L004 |
| AL320_153 | 2008 | 310 | M | 5 | 2.5175976 | 3 | 0.6264462774 | 1.666575662 | 2.269380261 | NA | NA | NA | NA | random | bin3 | 7-07-AL320-153_S7_L004 |
| AL320_263 | 2008 | 300 | M | 5 | 2.847006 | 3 | 0.4863638243 | 1.589579793 | 2.6 | NA | NA | NA | NA | random | bin3 | 1-01-AL320-263_S1_L004 |
| AL320_331 | 2008 | 430 | F | 6 | 3.670518 | 6 | 0.6941019 | 1.4470317 | 2.0705454 | 2.5058325 | 3.1528773 | 3.5 | NA | random | bin3 | 17-17-AL320-331_S17_L004 |
| AL320_335 | 2008 | 490 | M | 5 | 4.505796 | 6 | 0.447054 | 1.247034 | 2.0940774 | 2.9175948 | 3.7999287 | 4.3 | NA | random | bin3 | 31-31-AL320-335_S31_L004 |
| AL320_379 | 2008 | 350 | M | 5 | 3.599931 | 3 | 0.75 | 2.45 | 3.4 | NA | NA | NA | NA | random | bin3 | 6-06-AL320-379_S6_L004 |
| AL320_398 | 2008 | 420 | F | 5 | 3.576399 | 6 | 0.6963977649 | 1.192131415 | 1.794099079 | 2.431479987 | 2.903610626 | 3.328529286 | NA | random | bin3 | 44-44-AL320-398_S44_L004 |

| | | | | | | | | | | | | | | | | |
|---|---|---|---|---|---|---|---|---|---|---|---|---|---|---|---|---|
| AL320_487 | 2008 | 240 | F | 5 | 2.3881875 | 3 | 0.3294024 | 1.3293867 | 1.9646661 | NA | NA | NA | NA | random | bin3 | 41-41-AL320-487_S41_L004 |
| AL320_5 | 2008 | 370 | F | 5 | 3.235227 | 4 | 0.5058714 | 1.2235044 | 2.117601 | 3.03523188 | NA | NA | NA | random | bin3 | 35-35-AL320-5_S35_L004 |
| AL320_532 | 2008 | 390 | F | 5 | 3.094062 | 3 | 0.5646987 | 1.9881987 | 2.79995067 | NA | NA | NA | NA | random | bin3 | 9-09-AL320-532_S9_L004 |
| AL320_772 | 2008 | 430 | F | 5 | 3.505815 | 3 | 0.6514159446 | 2.226670653 | 3.375530361 | NA | NA | NA | NA | random | bin3 | 10-10-AL320-772_S10_L004 |
| AL320_865 | 2008 | 270 | M | 6 | 2.7293613 | 3 | 0.7541682573 | 1.831546336 | 2.5 | NA | NA | NA | NA | random | bin3 | 12-12-AL320-865_S12_L004 |
| AL320_879 | 2008 | 400 | F | 6 | 3.775266 | 4 | 0.494109 | 1.4940825 | 1.941141 | 3.1399809 | NA | NA | NA | random | bin3 | 52-52-AL320-879_S52_L004 |
| AL324_1047 | 2008 | 510 | M | 6 | 4.035219 | 5 | 0.6489411397 | 1.581052027 | 2.383374686 | 3.457073306 | 3.85 | NA | NA | random | bin3 | 103-103-AL324-1047_S49_L004 |
| AL324_1064 | 2008 | 440 | M | 6 | 3.870516 | 4 | 0.8048787446 | 1.941175432 | 2.923599148 | 3.468076549 | NA | NA | NA | random | bin3 | 74-74-AL324-1064_S20_L004 |
| AL324_1151 | 2008 | 480 | F | 8 | 4.105803 | 5 | 0.63528 | 1.8940794 | 2.5411272 | 3.2234691 | 3.8 | NA | NA | random | bin3 | 32-32-AL324-1151_S32_L004 |
| AL324_652 | 2008 | 220 | M | 6 | 2.2470147 | 2 | 0.7568868586 | 2.005097484 | NA | NA | NA | NA | NA | random | bin3 | 80-80-AL324-652_S26_L004 |
| AL435_298 | 2014 | 320 | F | 5 | 2.8940655 | 3 | 0.6286303753 | 2.348460953 | 2.8 | NA | NA | NA | NA | random | bin4 | J35428-S1-_L1_S125_L002 |
| AL435_334 | 2014 | 310 | M | 5 | 2.8705371 | 3 | 0.8065983966 | 1.991122238 | 2.8 | NA | NA | NA | NA | random | bin4 | J35413-S1-_L1_S110_L002 |
| AL435_403 | 2014 | 320 | M | 5 | 3.976395 | 5 | 0.7433609596 | 1.486723725 | 2.595865011 | 3.398224796 | 3.9 | NA | NA | random | bin4 | J35419-S1-_L1_S116_L002 |
| AL435_413 | 2014 | 270 | M | 6 | 2.741127 | 4 | 0.7594484835 | 1.756220117 | 2.183408262 | 2.7 | NA | NA | NA | random | bin4 | J35426-S1-_L1_S123_L002 |
| AL435_634 | 2014 | 390 | F | 5 | 3.023472 | 3 | 0.4742735811 | 2.07493539 | 3 | NA | NA | NA | NA | random | bin4 | J35414-S1-_L1_S111_L002 |
| AL435_636 | 2014 | 410 | M | 5 | 3.494052 | 5 | 0.7764564 | 1.7529093 | 2.2470129 | 2.8234758 | 3.4 | NA | NA | random | bin4 | J35427-S1-_L1_S124_L002 |
| AL435_721 | 2014 | 330 | F | 7 | 3.023472 | 4 | 0.8536846452 | 1.659944381 | 2.371347043 | 3 | NA | NA | NA | random | bin4 | J35424-S1-_L1_S121_L002 |
| AL435_725 | 2014 | 310 | M | 6 | 2.5646601 | 2 | 0.7293984 | 2.36466204 | NA | NA | NA | NA | NA | random | bin4 | J35434-S1-_L1_S131_L002 |
| AL435_730 | 2014 | 330 | M | 6 | 3.07053 | 3 | 0.6588114 | 1.141155 | 2.564661 | NA | NA | NA | NA | random | bin4 | J35423-S1-_L1_S120_L002 |
| AL437_245 | 2014 | 320 | M | 6 | 2.8352403 | 3 | 0.7088129519 | 1.937410822 | 2.646223894 | NA | NA | NA | NA | random | bin4 | J35406-S1-_L1_S103_L002 |
| AL437_251 | 2014 | 290 | M | 6 | 2.5646544 | 3 | 0.6470451 | 1.835256 | 2.39995323 | NA | NA | NA | NA | random | bin4 | J35395-S1-_L1_S92_L002 |
| AL437_264 | 2014 | 320 | M | 6 | 3.294054 | 4 | 0.5213653754 | 1.421896367 | 2.606803913 | 3.1 | NA | NA | NA | random | bin4 | J35407-S1-_L1_S104_L002 |

| | | | | | | | | | | | | | | | | |
|---|---|---|---|---|---|---|---|---|---|---|---|---|---|---|---|---|
| AL437_288 | 2014 | 350 | F | 5 | 3.129351 | 3 | 0.6588099 | 2.2234857 | 2.92935297 | NA | NA | NA | NA | random | bin4 | J35396-S1-_L1_S93_L002 |
| AL437_505 | 2014 | 230 | M | 6 | 2.1058419 | 3 | 0.6470508 | 1.2940959 | 1.92937512 | NA | NA | NA | NA | random | bin4 | J35398-S1-_L1_S95_L002 |
| AL437_562 | 2014 | 430 | F | 6 | 3.563544 | 4 | 0.5210818746 | 1.338224334 | 2.79376555 | 3.4 | NA | NA | NA | random | bin4 | J35425-S1-_L1_S122_L002 |
| AL437_573 | 2014 | 460 | F | 6 | 3.729339 | 5 | 0.793224895 | 1.988981835 | 2.770365723 | 3.1 | 3.5 | NA | NA | random | bin4 | J35399-S1-_L1_S96_L002 |
| AL437_579 | 2014 | 420 | F | 5 | 2.9528877 | 3 | 0.566954701 | 1.996152263 | 2.8 | NA | NA | NA | NA | random | bin4 | J35397-S1-_L1_S94_L002 |
| AL437_582 | 2014 | 320 | F | 5 | 3.223467 | 5 | 0.5214416408 | 1.113990801 | 1.730242567 | 2.405749341 | 3 | NA | NA | random | bin4 | J35432-S1-_L1_S129_L002 |
| AL437_601 | 2014 | 340 | F | 6 | 3.446994 | 5 | 0.6278031271 | 1.445134023 | 2.262461899 | 2.736272713 | 3.4 | NA | NA | random | bin4 | J35405-S1-_L1_S102_L002 |
| AL437_611 | 2014 | 310 | F | 5 | 3.282297 | 4 | 0.7764591 | 1.9764387 | 2.4940734 | 3.1 | NA | NA | NA | random | bin4 | J35433-S1-_L1_S130_L002 |
| AL521_566 | 2019 | 300 | M | 5 | 2.3999532 | 3 | 0.7684568129 | 1.761544264 | 2.2 | NA | NA | NA | NA | random | bin5 | J35415-S1-_L1_S112_L002 |
| AL521_726 | 2019 | 280 | M | 5 | 2.7764181 | 5 | 0.5339250681 | 0.8542825901 | 1.649239937 | 2.230626713 | 2.7 | NA | NA | random | bin5 | J35416-S1-_L1_S113_L002 |
| AL521_847 | 2019 | 380 | M | 5 | 3.152883 | 5 | 0.5058723 | 1.0352757 | 1.72938 | 2.541129 | 2.92935663 | NA | NA | random | bin5 | J35420-S1-_L1_S117_L002 |
| AL521_865 | 2019 | 270 | M | 5 | 2.6587704 | 3 | 0.6381021358 | 2.15065042 | 2.6 | NA | NA | NA | NA | random | bin5 | J35421-S1-_L1_S118_L002 |
| AL522_473 | 2019 | 230 | M | 6 | 1.9785411 | 2 | 0.6056997595 | 1.836523718 | NA | NA | NA | NA | NA | random | bin5 | J35403-S1-_L1_S100_L002 |
| AL522_510 | 2019 | 290 | F | 5 | 2.5646601 | 3 | 0.6263930347 | 1.938266975 | 2.3 | NA | NA | NA | NA | random | bin5 | J35418-S1-_L1_S115_L002 |
| AL522_511 | 2019 | 250 | M | 6 | 2.741127 | 4 | 0.7148181803 | 1.199241196 | 2.27442566 | 2.7 | NA | NA | NA | random | bin5 | J35404-S1-_L1_S101_L002 |
| AL522_519 | 2019 | 360 | M | 7 | 3.22347 | 5 | 0.7999857 | 1.8705519 | 2.4234837 | 2.811714 | 3.2 | NA | NA | random | bin5 | J35400-S1-_L1_S97_L002 |
| AL522_531 | 2019 | 330 | F | 5 | 2.9293536 | 4 | 0.8386457811 | 1.535552474 | 2.374197954 | 2.85 | NA | NA | NA | random | bin5 | J35431-S1-_L1_S128_L002 |
| AL522_532 | 2019 | 280 | F | 5 | 2.6705391 | 3 | 0.6617258696 | 2.06789564 | 2.469656008 | NA | NA | NA | NA | random | bin5 | J35411-S1-_L1_S108_L002 |
| AL522_543 | 2019 | 220 | M | 6 | 2.1764298 | 2 | 0.5708696617 | 1.926680718 | NA | NA | NA | NA | NA | random | bin5 | J35401-S1-_L1_S98_L002 |
| AL522_547 | 2019 | 370 | F | 7 | 2.8352403 | 4 | 0.6761882658 | 1.565906436 | 2.277681154 | 2.8 | NA | NA | NA | random | bin5 | J35402-S1-_L1_S99_L002 |
| AL522_561 | 2019 | 300 | F | 5 | 2.43525 | 3 | 0.6029011425 | 1.66684797 | 2.246103743 | NA | NA | NA | NA | random | bin5 | J35408-S1-_L1_S105_L002 |
| AL522_564 | 2019 | 300 | M | 6 | 3.011706 | 5 | 0.6639975875 | 1.470283612 | 2.098707352 | 2.572997001 | 2.9 | NA | NA | random | bin5 | J35409-S1-_L1_S106_L002 |

| | | | | | | | | | | | | | | | | |
|---|---|---|---|---|---|---|---|---|---|---|---|---|---|---|---|---|
| AL522_579 | 2019 | 320 | F | 5 | 2.8234746 | 3 | 0.7646895 | 2.0117226 | 2.62347654 | NA | NA | NA | NA | random | bin5 | J35412-S1-_L1_S109_L002 |
| AL522_593 | 2019 | 330 | F | 7 | 2.6929806 | 3 | 0.5207789457 | 1.087076907 | 2.113984798 | NA | NA | NA | NA | random | bin5 | J35422-S1-_L1_S119_L002 |
| AL522_594 | 2019 | 300 | F | 5 | 2.7646524 | 3 | 0.5576724108 | 1.95780449 | 2.645997957 | NA | NA | NA | NA | random | bin5 | J35429-S1-_L1_S126_L002 |
| AL522_613 | 2019 | 270 | M | 5 | 2.3646681 | 3 | 0.8823399 | 1.9176096 | 2.18819544 | NA | NA | NA | NA | random | bin5 | J35410-S1-_L1_S107_L002 |
| AL522_624 | 2019 | 300 | F | 5 | 2.8352343 | 5 | 0.4370949496 | 1.051398175 | 1.96103226 | 2.350880647 | 2.8 | NA | NA | random | bin5 | J35430-S1-_L1_S127_L002 |
| AL522_682 | 2019 | 410 | F | 5 | 3.199944 | 4 | 0.4235259 | 0.9764517 | 2.4705456 | 3 | NA | NA | NA | random | bin5 | J35417-S1-_L1_S114_L002 |

**Table S2. Estimated von Bertalanffy growth parameters for individual fish, catch years and all samples.**

The Bayesian hierarchical model estimates parameters of nested levels at the same time. Here, three estimated von Bertalanffy parameters L∞, length at infinity, k, a growth coefficient, and t0, hypothetical length at age 0, were estimated for all samples, for each bin, based on the catch years, and for each individual. The individual parameters were used to calculate growth performance, Φ, which was used in genotype-phenotype association analysis.

| | **L∞** | **k** | **t0** |
|---|---|---|---|
| All | 7.4265107 | 0.1785009186 | 0.45453227 |
| **group parameters** | | | |
| bin1 | 9.197300231 | 0.128748689 | 0.4674114129 |
| bin2 | 8.07270249 | 0.1346183828 | 0.4368796201 |
| bin3 | 8.355492139 | 0.1339595874 | 0.4697016901 |
| bin4 | 5.601883356 | 0.2381987686 | 0.4521721634 |
| bin5 | 4.794222888 | 0.2540789252 | 0.4417218798 |
| **Individual parameters** | | | |
| AL097_334 | 9.450599562 | 0.1520235359 | 0.4475552977 |
| AL097_430 | 9.087302412 | 0.1130445062 | 0.4992249003 |
| AL097_552 | 8.991693021 | 0.1089703578 | 0.4471818953 |
| AL099_189 | 8.87386943 | 0.1081162503 | 0.4034060656 |
| AL099_326 | 9.001873984 | 0.1064093858 | 0.4727587402 |
| AL099_328 | 9.294134062 | 0.1440551036 | 0.458454618 |
| AL099_344 | 9.107911539 | 0.1144948398 | 0.5125570973 |
| AL099_35 | 9.862030975 | 0.1545818366 | 0.6151723101 |
| AL099_37 | 9.338815522 | 0.179830962 | 0.4050394258 |
| AL099_577 | 9.886988158 | 0.1854323845 | 0.4736466419 |
| AL099_591 | 8.922809483 | 0.08965639297 | 0.4861141298 |
| AL099_597 | 9.519168066 | 0.1661813843 | 0.388586572 |
| AL099_600 | 9.04811865 | 0.1077003178 | 0.5133851482 |
| AL099_602 | 8.865169484 | 0.08686046859 | 0.4291770037 |
| AL099_613 | 9.703720402 | 0.1723847808 | 0.4776966284 |
| AL099_621 | 9.153721997 | 0.1159366327 | 0.5329026519 |
| AL099_630 | 9.179612099 | 0.1220113227 | 0.510800712 |
| AL099_636 | 9.269059273 | 0.1181639864 | 0.5105184667 |
| AL099_78 | 9.017167028 | 0.09577655554 | 0.5042855393 |
| AL101_105 | 9.300229768 | 0.155219942 | 0.4468083347 |
| AL101_114 | 9.174572274 | 0.1124477676 | 0.4484334979 |
| AL101_185 | 9.131668638 | 0.1103295181 | 0.5051449708 |
| AL101_214 | 9.07790325 | 0.1186426579 | 0.4450988971 |
| AL101_287 | 9.339071435 | 0.1272314452 | 0.5661345765 |
| AL101_288 | 8.941416968 | 0.09725037072 | 0.4703341392 |
| AL101_290 | 9.408169079 | 0.1379044579 | 0.5625363926 |

| | | | |
|---|---|---|---|
| AL101_332 | 9.081519588 | 0.1115161007 | 0.5328694771 |
| AL101_344 | 9.221909288 | 0.1375316857 | 0.377767302 |
| AL101_422 | 9.036769493 | 0.1108161478 | 0.4243746867 |
| AL101_423 | 9.507684883 | 0.154410608 | 0.4506158986 |
| AL101_430 | 9.042936003 | 0.09719637536 | 0.4990055604 |
| AL101_434 | 8.737914906 | 0.07886077014 | 0.4409890429 |
| AL101_442 | 9.086019789 | 0.134018513 | 0.423991964 |
| AL101_504 | 9.271457586 | 0.1230483967 | 0.5372382919 |
| AL101_523 | 9.441536901 | 0.1366415583 | 0.5338619759 |
| AL101_546 | 9.17519508 | 0.1277895548 | 0.4689240648 |
| AL101_595 | 9.14239765 | 0.1131475528 | 0.5551555344 |
| AL101_623 | 8.956316965 | 0.09746956772 | 0.4959048311 |
| AL101_679 | 9.043583385 | 0.1060935329 | 0.5175851339 |
| AL111_235 | 9.461991327 | 0.15358678 | 0.4465150743 |
| AL111_28 | 9.07518508 | 0.1083889961 | 0.5008817759 |
| AL114_826 | 9.526137235 | 0.1609316128 | 0.4221965564 |
| AL114_868 | 9.475413075 | 0.1582214552 | 0.4294103049 |
| AL114_937 | 9.532596612 | 0.1626247699 | 0.4209297711 |
| AL129_106 | 9.002026372 | 0.1097054651 | 0.4810656196 |
| AL129_148 | 8.794570505 | 0.07754371282 | 0.4536293287 |
| AL129_243 | 9.245132909 | 0.1332643984 | 0.5029691637 |
| AL129_266 | 8.950702232 | 0.09715883083 | 0.4819860694 |
| AL129_339 | 9.109785129 | 0.1231829907 | 0.4162388817 |
| AL129_369 | 9.09230404 | 0.1182275187 | 0.4620745171 |
| AL129_377 | 8.851164347 | 0.08940231367 | 0.4486286454 |
| AL129_389 | 9.416035586 | 0.1482700369 | 0.5093214217 |
| AL129_401 | 9.662875585 | 0.1686834452 | 0.4689490516 |
| AL129_419 | 9.040034327 | 0.1117776739 | 0.4653045182 |
| AL129_467 | 9.010387282 | 0.1080124261 | 0.4631736665 |
| AL129_47 | 9.002389964 | 0.1044357431 | 0.4646519995 |
| AL129_87 | 9.591999296 | 0.1638570041 | 0.4284949264 |
| AL130_199 | 9.058881759 | 0.1082911369 | 0.4867830913 |
| AL130_206 | 9.47053881 | 0.1548214306 | 0.4071403752 |
| AL130_223 | 9.374073623 | 0.1471718248 | 0.4393894005 |
| AL130_244 | 9.408045541 | 0.1498888662 | 0.4189800309 |
| AL130_429 | 9.406646004 | 0.1473586536 | 0.4953509983 |
| AL130_440 | 9.583337471 | 0.1582945219 | 0.4978265952 |
| AL130_68 | 9.249965364 | 0.1408448746 | 0.4081199343 |
| AL133_100 | 9.49050287 | 0.1579455583 | 0.4210616173 |
| AL133_116 | 9.200936866 | 0.1280703676 | 0.4736767546 |
| AL133_119 | 9.163756832 | 0.1232225343 | 0.4880580957 |
| AL133_136 | 9.117893336 | 0.1259574667 | 0.4129214144 |
| AL207_131 | 7.649680902 | 0.1081416354 | 0.4202711597 |
| AL207_143 | 7.795216507 | 0.1114914491 | 0.4674499961 |

| | | | |
|---|---|---|---|
| AL207_146 | 8.161805783 | 0.138122074 | 0.4347289713 |
| AL207_149 | 7.89575326 | 0.105939195 | 0.4810880663 |
| AL207_151 | 8.598745009 | 0.1671428625 | 0.4109410267 |
| AL207_197 | 8.306722064 | 0.1450613559 | 0.4501820786 |
| AL207_20 | 8.02409794 | 0.1322939132 | 0.4251877645 |
| AL207_231 | 8.245481996 | 0.1461482888 | 0.4120184535 |
| AL207_249 | 8.29965245 | 0.1428372343 | 0.4499806396 |
| AL207_25 | 8.297817071 | 0.1490380849 | 0.4018941439 |
| AL207_413 | 8.011895365 | 0.1381896123 | 0.3946931393 |
| AL207_462 | 8.177615271 | 0.1419191474 | 0.4198506593 |
| AL207_473 | 7.936805647 | 0.1243798414 | 0.4476587427 |
| AL207_490 | 7.984126882 | 0.1316682425 | 0.3958273588 |
| AL207_504 | 7.721832776 | 0.1101414055 | 0.4334560669 |
| AL207_516 | 8.026537311 | 0.1345181681 | 0.4081450666 |
| AL207_539 | 7.871298514 | 0.1167095049 | 0.4777288743 |
| AL207_561 | 8.267333165 | 0.1404821668 | 0.4525514602 |
| AL207_61 | 8.045089124 | 0.1303392316 | 0.4531575304 |
| AL207_625 | 8.516871434 | 0.1609191553 | 0.4573959705 |
| AL207_691 | 8.375366546 | 0.1543463303 | 0.3911525867 |
| AL207_91 | 7.652973621 | 0.1049047922 | 0.4250222648 |
| AL318_200 | 8.004324218 | 0.1055665781 | 0.6565550876 |
| AL318_234 | 7.713886643 | 0.09961048103 | 0.4345559428 |
| AL318_29 | 8.296078913 | 0.1150727245 | 0.7374793351 |
| AL318_9 | 7.904237062 | 0.104625754 | 0.5302913157 |
| AL320_1003 | 8.518634158 | 0.133939944 | 0.6414515207 |
| AL320_1067 | 8.470923633 | 0.1378787445 | 0.3915099997 |
| AL320_1172 | 8.534894472 | 0.1434299212 | 0.3859437223 |
| AL320_132 | 8.582672465 | 0.1386076274 | 0.6260217212 |
| AL320_153 | 8.291394134 | 0.1303608222 | 0.4312805614 |
| AL320_263 | 8.546185505 | 0.1403667817 | 0.5117499617 |
| AL320_331 | 7.763856655 | 0.1113906249 | 0.3479169119 |
| AL320_335 | 8.449163894 | 0.1279784037 | 0.5855312028 |
| AL320_379 | 9.183784065 | 0.1743276474 | 0.4148729795 |
| AL320_398 | 7.755807895 | 0.1029128544 | 0.3921064958 |
| AL320_487 | 8.090126178 | 0.11787785 | 0.5356935788 |
| AL320_5 | 8.270666667 | 0.125019782 | 0.527420293 |
| AL320_532 | 8.739423257 | 0.1513249592 | 0.4638133059 |
| AL320_772 | 9.15779552 | 0.1719636176 | 0.4647211654 |
| AL320_865 | 8.446664257 | 0.1387328548 | 0.3884157674 |
| AL320_879 | 8.265953535 | 0.126381176 | 0.505030336 |
| AL324_1047 | 8.444693844 | 0.1370481296 | 0.4576930795 |
| AL324_1064 | 8.589451741 | 0.1481852295 | 0.3686268299 |
| AL324_1151 | 8.32059804 | 0.1364180702 | 0.3853341051 |
| AL324_652 | 8.731817382 | 0.1512966595 | 0.4038428104 |

| | | | |
|---|---|---|---|
| AL435_298 | 5.864809261 | 0.2652378212 | 0.4475483427 |
| AL435_334 | 5.787893591 | 0.2559622744 | 0.4412232761 |
| AL435_403 | 5.749631241 | 0.2394476233 | 0.4614281932 |
| AL435_413 | 5.29856728 | 0.2187984678 | 0.4363076765 |
| AL435_634 | 5.91046528 | 0.2627780346 | 0.467530867 |
| AL435_636 | 5.352833388 | 0.2234446852 | 0.437419918 |
| AL435_721 | 5.453864234 | 0.229488045 | 0.4387939227 |
| AL435_725 | 6.01337452 | 0.2734623517 | 0.4443419378 |
| AL435_730 | 5.47145833 | 0.2201121101 | 0.4649564816 |
| AL437_245 | 5.699779538 | 0.2478242769 | 0.4472282347 |
| AL437_251 | 5.547323367 | 0.2348889923 | 0.4491953529 |
| AL437_264 | 5.561382498 | 0.229835008 | 0.469202689 |
| AL437_288 | 5.9140313 | 0.2658132443 | 0.4494727246 |
| AL437_505 | 5.214409517 | 0.1973634758 | 0.453997777 |
| AL437_562 | 5.736571089 | 0.2388926366 | 0.478075704 |
| AL437_573 | 5.475800125 | 0.2460417728 | 0.4290892014 |
| AL437_579 | 5.789128394 | 0.254111154 | 0.4596674789 |
| AL437_582 | 5.161057285 | 0.1825767766 | 0.4649460912 |
| AL437_601 | 5.365783609 | 0.2146789286 | 0.4558613664 |
| AL437_611 | 5.551650309 | 0.2405733583 | 0.436009765 |
| AL521_566 | 4.785374419 | 0.2576090993 | 0.4200144405 |
| AL521_726 | 4.599358919 | 0.187787258 | 0.4622705359 |
| AL521_847 | 4.673541333 | 0.2088844679 | 0.4659267746 |
| AL521_865 | 4.931469905 | 0.2952174229 | 0.4286036625 |
| AL522_473 | 4.860981505 | 0.2732937974 | 0.4394298411 |
| AL522_510 | 4.828798825 | 0.2692200531 | 0.4311532453 |
| AL522_511 | 4.739056117 | 0.2372747965 | 0.4437710032 |
| AL522_519 | 4.737025053 | 0.2642862543 | 0.4062924007 |
| AL522_531 | 4.780649417 | 0.2570631968 | 0.4215520795 |
| AL522_532 | 4.885994957 | 0.2850281813 | 0.4258320717 |
| AL522_543 | 4.886445622 | 0.2783876422 | 0.4406260105 |
| AL522_547 | 4.763554709 | 0.2521552246 | 0.436198989 |
| AL522_561 | 4.783560822 | 0.2544689779 | 0.4399686355 |
| AL522_564 | 4.647066793 | 0.229708281 | 0.4300718671 |
| AL522_579 | 4.912403487 | 0.2922403256 | 0.4199102108 |
| AL522_593 | 4.693241561 | 0.2201993833 | 0.4616249098 |
| AL522_594 | 4.923511456 | 0.2888315373 | 0.4431665049 |
| AL522_613 | 4.792925566 | 0.2659231351 | 0.4057731667 |
| AL522_624 | 4.62913837 | 0.2064427007 | 0.4635240498 |
| AL522_682 | 4.845561151 | 0.2483105756 | 0.4853192219 |

**Table S3. A list of enriched GO terms for outlier windows $F_{st}$ between 1996 and 2019.**

$F_{st}$ values in non-overlapping 50Kb windows along the genome were calculated for 1996 and 2019 to identify regions that are the most differentiated over time. The windows with highest 5% $F_{st}$ were assigned as outlier windows. The genes residing in these outlier windows were subjected to gene ontology (GO) enrichment test to identify any biological functions that have differentiated over time. P values are adjusted using false discovery rate. Only biological processes were presented among GO categories for the analysis.

| GO.term | GO.name | p.value.adjusted |
|---|---|---|
| GO:0006468 | protein phosphorylation | 0.02754635203 |
| GO:0006541 | glutamine metabolic process | 0.02797620132 |
| GO:0019217 | regulation of fatty acid metabolic process | 0.02797620132 |
| GO:0016079 | synaptic vesicle exocytosis | 0.02797620132 |
| GO:0031998 | regulation of fatty acid beta-oxidation | 0.02797620132 |
| GO:1990504 | dense core granule exocytosis | 0.02797620132 |
| GO:0006171 | cAMP biosynthetic process | 0.02797620132 |
| GO:0034220 | ion transmembrane transport | 0.02870276413 |
| GO:0019395 | fatty acid oxidation | 0.04186881996 |
| GO:0042157 | lipoprotein metabolic process | 0.04212093394 |
| GO:0051301 | cell division | 0.04212093394 |
| GO:0061074 | regulation of neural retina development | 0.04212093394 |
| GO:0033077 | T cell differentiation in thymus | 0.04212093394 |
| GO:0006044 | N-acetylglucosamine metabolic process | 0.04212093394 |
| GO:0006432 | phenylalanyl-tRNA aminoacylation | 0.04212093394 |
| GO:0070509 | calcium ion import | 0.04212093394 |
| GO:0062012 | regulation of small molecule metabolic process | 0.04212093394 |
| GO:0007166 | cell surface receptor signaling pathway | 0.04212093394 |
| GO:0030178 | negative regulation of Wnt signaling pathway | 0.04212093394 |
| GO:0006814 | sodium ion transport | 0.04212093394 |
| GO:0005975 | carbohydrate metabolic process | 0.04212093394 |
| GO:0007264 | small GTPase mediated signal transduction | 0.04212093394 |
| GO:0140013 | meiotic nuclear division | 0.04212093394 |
| GO:0007169 | transmembrane receptor protein tyrosine kinase signaling pathway | 0.04212093394 |
| GO:0009190 | cyclic nucleotide biosynthetic process | 0.04212093394 |
| GO:0060027 | convergent extension involved in gastrulation | 0.04212093394 |
| GO:0006796 | phosphate-containing compound metabolic process | 0.04212093394 |
| GO:0030968 | endoplasmic reticulum unfolded protein response | 0.04212093394 |
| GO:0030098 | lymphocyte differentiation | 0.04212093394 |
| GO:0061982 | meiosis I cell cycle process | 0.04212093394 |
| GO:0001702 | gastrulation with mouth forming second | 0.04212093394 |
| GO:0050794 | regulation of cellular process | 0.04212093394 |
| GO:0010033 | response to organic substance | 0.04212093394 |
| GO:0009081 | branched-chain amino acid metabolic process | 0.04212093394 |
| GO:0007099 | centriole replication | 0.04212093394 |

| | | |
|---|---|---|
| GO:0035567 | non-canonical Wnt signaling pathway | 0.04212093394 |
| GO:0032007 | negative regulation of TOR signaling | 0.04212093394 |
| GO:0021952 | central nervous system projection neuron axonogenesis | 0.04212093394 |
| GO:0055074 | calcium ion homeostasis | 0.04212093394 |
| GO:0009062 | fatty acid catabolic process | 0.04212093394 |
| GO:0050769 | positive regulation of neurogenesis | 0.04212093394 |
| GO:0042110 | T cell activation | 0.04212093394 |
| GO:0035967 | cellular response to topologically incorrect protein | 0.04212093394 |
| GO:0030433 | ubiquitin-dependent ERAD pathway | 0.04212093394 |
| GO:0060028 | convergent extension involved in axis elongation | 0.04212093394 |
| GO:0016579 | protein deubiquitination | 0.04285964063 |
| GO:0030001 | metal ion transport | 0.04285964063 |
| GO:0007154 | cell communication | 0.04331273079 |
| GO:0048729 | tissue morphogenesis | 0.04501724582 |
| GO:0048585 | negative regulation of response to stimulus | 0.04538697515 |
| GO:0009247 | glycolipid biosynthetic process | 0.04719048324 |
| GO:0023052 | signaling | 0.04719048324 |
| GO:0065004 | protein-DNA complex assembly | 0.04790241745 |
| GO:0090162 | establishment of epithelial cell polarity | 0.04790241745 |
| GO:0035825 | homologous recombination | 0.04790241745 |
| GO:0007131 | reciprocal meiotic recombination | 0.04790241745 |
| GO:0006040 | amino sugar metabolic process | 0.04790241745 |
| GO:0006801 | superoxide metabolic process | 0.04790241745 |
| GO:0098655 | cation transmembrane transport | 0.04790241745 |
| GO:0048285 | organelle fission | 0.04790241745 |
| GO:0023057 | negative regulation of signaling | 0.04790241745 |
| GO:0010648 | negative regulation of cell communication | 0.04790241745 |
| GO:0051716 | cellular response to stimulus | 0.04790241745 |
| GO:0023061 | signal release | 0.04790241745 |
| GO:0032402 | melanosome transport | 0.04790241745 |
| GO:0006284 | base-excision repair | 0.04790241745 |
| GO:0036211 | protein modification process | 0.04790241745 |
| GO:0043632 | modification-dependent macromolecule catabolic process | 0.04790241745 |
| GO:0006511 | ubiquitin-dependent protein catabolic process | 0.04790241745 |
